# A potent alpaca-derived nanobody that neutralizes SARS-CoV-2 variants

**DOI:** 10.1101/2022.01.18.476801

**Authors:** Jules B. Weinstein, Timothy A. Bates, Hans C. Leier, Savannah K. McBride, Eric Barklis, Fikadu G. Tafesse

## Abstract

The spike glycoprotein of SARS-CoV-2 engages with human angiotensin-converting enzyme 2 (ACE2) to facilitate infection. Here, we describe an alpaca-derived heavy chain antibody fragment (VHH), saRBD-1, that disrupts this interaction by competitively binding to the spike protein receptor-binding domain. We further generated an engineered bivalent nanobody construct engineered by a flexible linker, and a dimeric Fc conjugated nanobody construct. Both multivalent nanobodies blocked infection at picomolar concentrations and demonstrated no loss of potency against emerging variants of concern including Alpha (B.1.1.7), Beta (B.1.351), Gamma (P.1), Epsilon (B.1.427/429), and Delta (B.1.617.2). saRBD-1 tolerates elevated temperature, freeze-drying, and nebulization, making it an excellent candidate for further development into a therapeutic approach for COVID-19.

## Introduction

The COVID-19 pandemic, caused by severe acute respiratory syndrome coronavirus-2 (SARS-CoV-2), is an ongoing global health crisis with over 230 million cases, 4.8 million deaths world-wide as of October 2021 (Dong et al., 2020). While several effective vaccines have been developed, concern about potential future surges of infections remain, due to the proliferation and spread of multiple variant strains, combined with waning protection from vaccination (Levin et al., 2021; Shrotri et al., 2021). It is anticipated that additional variants will continue to emerge, and the slow pace of global vaccination creates greater opportunity for emergence and spread of vaccine resistant variants (Luo et al., 2021).

SARS-CoV-2 is an enveloped, positive-sense, single-stranded RNA virus, and a member of the Coronaviridae family, so named for the crown-like protrusions visible on their outer membranes in EM micrographs (Huang et al., 2020). Four structural proteins are encoded by SARS-CoV-2: spike (S), envelope, membrane, and nucleocapsid (Jiang et al., 2020). Homotrimers of the S glycoprotein form the characteristic crown-like protrusions on the virion surface, where it facilitates entry into cells through its interaction with the cell surface protein angiotensin-converting enzyme 2 (ACE2) (Hoffmann et al., 2020). Each monomer of S is composed of two subunits, S1 and S2, the former being responsible for ACE2 binding, and the latter involved in membrane fusion with target cells. These subunits are connected by a polybasic cleavage site, which is typically cleaved by the human cell surface-bound protease, TMPRSS2, releasing the S1 subunit to reveal the fusion peptide of S2 (Hoffmann et al., 2020). Many neutralizing antibodies function by binding to the receptor binding domain (RBD) of the S1 subunit, thereby blocking ACE2 engagement and preventing protease activation of fusion-competent S2 (Carrillo et al., 2021).

Neutralizing antibodies have been shown to be protective against COVID-19 disease (Khoury et al., 2021), and a majority of the treatment options approved for emergency use by the United States Food and Drug Administration for severe COVID-19 consist of monoclonal antibody cocktails (Kumar et al., 2021). The advantage of monoclonal antibodies is their ability to prevent entry of the virus into cells through their highly specific interaction with the spike protein (Jiang et al., 2020). This effectively limits the ability of SARS-CoV-2 to infect cells with minimal risk of side effects (Weinreich et al., 2021). The disadvantages of monoclonal antibody treatments are the difficulties of their production, high cost, and the possibility of escape by variants.

Several SARS-CoV-2 variants have displayed a propensity for increased transmission, as well as evasion of antibody neutralization by immune sera. The most clinically important of these are the variants of concern (VOC) including Alpha (B.1.1.7) (Bates et al., 2021a; Liu et al., 2021a; Planas et al., 2021), Beta (B.1.351) (Bates et al., 2021a), Gamma (P.1) (Bates et al., 2021b; Hoffmann et al., 2021), Delta (B.1.617.2) (Liu et al., 2021b), Epsilon (B.1.427/429) (Deng et al., 2021) and Omicron (B.1.529) (Liu et al., 2021c; Zhang et al., 2021); each of which has demonstrated significant immune evasion. These variants all incorporate numerous amino acid substitutions that are responsible for altering the epitopes critical for antibody-based neutralization. Previous work has shown that antibody cross-reactivity is common between different coronaviruses (Yuan et al., 2020). However, cross-neutralization is rare, and while some cross-neutralizing antibodies have been described (Pinto et al., 2020), even strongly binding cross-reactive antibodies are not necessarily neutralizing (Bates et al., 2021c).

One promising new technology that overcomes some of the inherent disadvantages of traditional monoclonal antibodies are nanobodies, which are immune fragments derived from the unique heavy-chain-only antibodies found in camelid species such as alpacas (Ingram et al., 2018). Composed solely of the heavy-chain only antibody variable domains (VHH), nanobodies are one-tenth the size of conventional antibodies, while preserving their binding affinities (Ingram et al., 2018). As single peptides with no need for glycosylation or complex maturation pathways, nanobodies offer several key advantages such as higher throughput discovery, simplified production, and improved stability. Because they lack antibody constant domains, nanobodies also avoid Fc-mediated immune activation (Salvador et al., 2019).

In this report, we detail the development of an alpaca-derived anti-SARS-CoV-2 nanobody (saRBD-1) with picomolar binding to the RBD portion of the spike protein. saRBD-1 displays high thermostability and remains functional after nebulization. Unmodified monovalent saRBD-1, bivalent saRBD-1, and a bivalent IgG Fc conjugated saRBD-1 protein all successfully neutralize live SARS-CoV-2 clinical isolates including the VOCs Alpha, Beta, Gamma, Epsilon, and Delta with no loss of potency. Variant cross-reactive nanobodies such as saRBD-1 may have therapeutic potential in COVID-19 caused by SARS-CoV-2 variants.

## Results

### A dominant VHH clone that binds SARS-COV-2 spike, saRBD-1 was isolated from spike RBD immunized alpaca

To acquire potent SARS-CoV-2 neutralizing VHHs, we first immunized an alpaca with purified SARS-CoV-2 S RBD. We used standard immunization techniques (Maass et al., 2007) over a 50-day immunization schedule, after which we generated a VHH gene library from immunized alpaca peripheral blood mononuclear cells (PBMCs), from which we isolated S binding VHH genes via phage display (Figure 1A). We performed panning against purified full-length trimeric S protein to maximize the number of native epitopes that match those present on live SARS-CoV-2 virus. Two rounds of panning enriched high binders in our library population. High quality hits were identified by high-throughput enzyme-linked immunosorbent assays (ELISA) of individual VHH clones on immobilized RBD. VHH hits which showed binding significantly above background were sequenced to determine their unique complementary-determining region 3 (CDR3) loops. Resulting high binding VHH sequence families with an enrichment of 10% or more after panning were tested for neutralizing activities. Neutralization assays used a GFP-reporter lentivirus pseudotyped with SARS-CoV-2 S protein (Crawford et al., 2020). Human ACE2 over-expressing HEK-293T cells (293T-ACE2) were incubated with pseudotyped virus in the presence of candidate VHHs. From our initial candidate pool, we discovered one novel VHH clone, referred to here as saRBD-1, that completely neutralized spike-mediated lentivirus transduction (Figure S1). We next analyzed the ability of saRBD-1 to associate with SARS-CoV-2 S using flow cytometry and immunofluorescence. African green monkey kidney cells (Vero E6) cells were infected with live SARS-CoV-2 WA1/2020 strain, then stained with anti-dsRNA monoclonal antibody to identify infected cells, and saRBD-1 or a control VHH (VHH52) (Cavallari, 2017) (Figure 1B). Cells positive for SARS-CoV-2 dsRNA showed concomitant binding by saRBD-1, but not VHH52 control. In a thermal shift assay, we found that equimolar saRBD-1 stabilized RBD protein and shifted the melting point by 8°C, from 52°C to 60°C (Figure S2). From this assay, we also determined that the melting point of saRBD-1 is 72°C in plain phosphate buffered saline (PBS) without stabilizing additives, indicating that it is highly stable. To corroborate the binding results with flow cytometry, S-transfected cells stained with saRBD1 and AlexaFlor488-anti-VHH antibody were 30% VHH-positive by our gating scheme, while control VHH52 treated cells and un-transfected control cells were VHH-negative (Figure 1C). Together, our data indicate that saRBD-1 binds strongly to native SARS-CoV-2 S protein.

**Figure 1:**
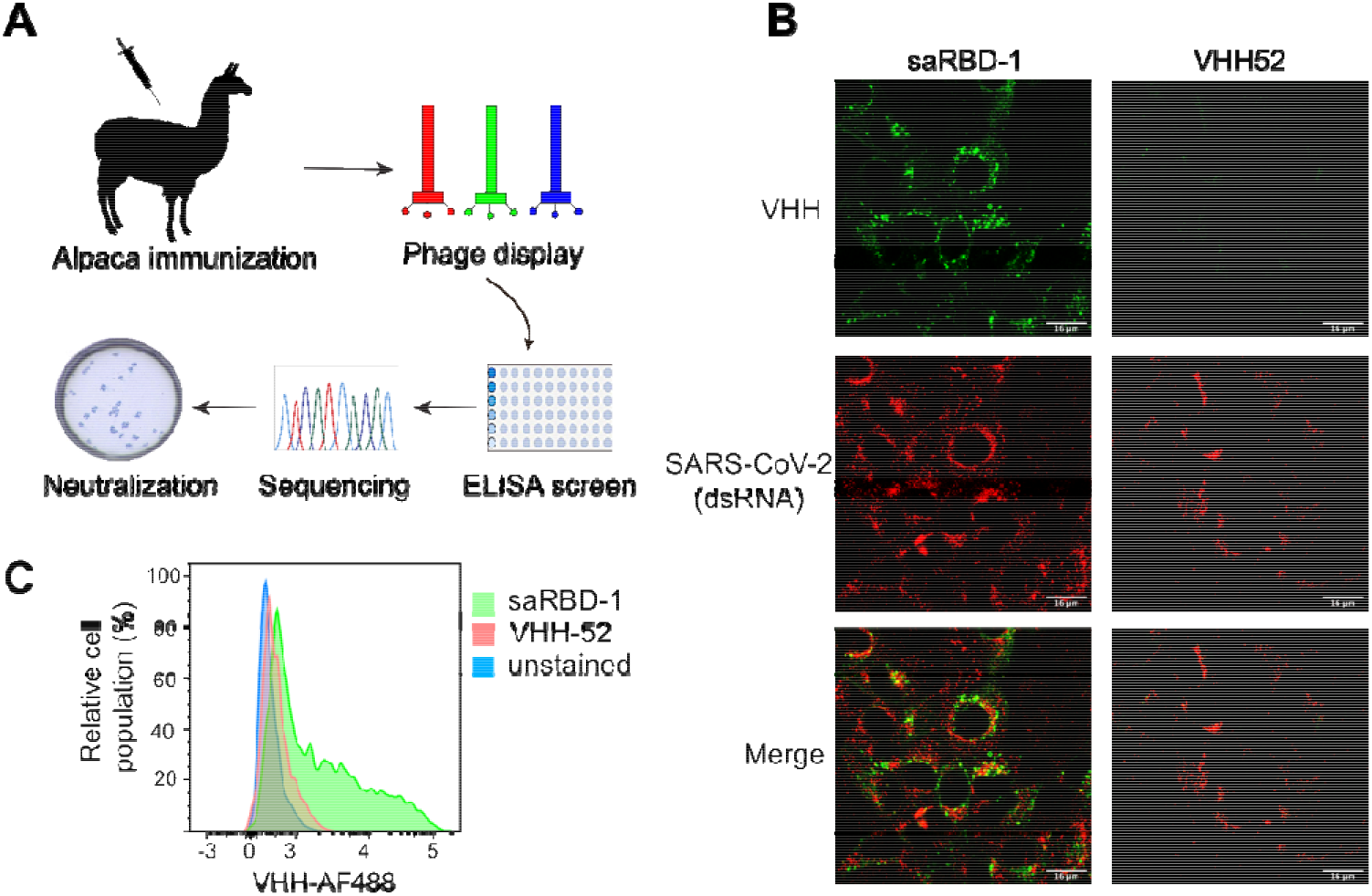
A dominant VHH clone that binds SARS-CoV-2 spike, saRBD-1 was isolated from an alpaca immunized with RBD. A) Schematic illustration of the immunization and VHH library construction pipeline. Alpacas were immunized over an 8-week period after which PBMC mRNA was isolated and processed into a VHH gene library. This library was transformed into phage-competent bacteria to generate a bacteriophage library, which was panned against SARS-CoV-2 S to enrich for binding clones. Clones were characterized through ELISAs on RBD and preliminary neutralization of S-pseudotyped lentivirus. B) Representative immunofluorescence staining showing saRBD-1 VHH specifically associates with SARS-CoV-2 infected VeroE6 cells. Cells were infected with SARS-CoV-2 virus for 24 hours. Fixed cells were stained with either saRBD-1 or control VHH (VHH52) followed by anti-VHH secondary (green). SARS-CoV-2 infection indicated by anti-dsRNA which stains replication centers (red). C) SARS-CoV-2 S-transfected cells are specifically bound by saRBD-1 at levels detectable by flow cytometry. 293T cells transfected with full-length SARS-CoV-2 S were stained with either VHH saRBD-1 (green) or VHH52 (red) control.

### VHH saRBD-1 binds SARS-CoV-2 spike and receptor domain with high affinity

We determined the subunit specificity of saRBD-1 by ELISA on purified full-length trimeric S, S1 (residues 14-684), RBD (residues 319-541), and S2 (residues 685-1273) proteins (Figure 2A). We found that saRBD-1 bound to full-length trimer with a 50% maximal binding response (EC_50_) of 100 pM, to S1 with an EC_50_ of 200 pM, and to RBD with an EC50 of 607 pM, while S2 showed no detectable binding, demonstrating that saRBD-1 binds specifically to the RBD subunit of the S protein (Figure 2B, D). Due to the promising initial binding characteristics of saRBD-1, we next investigated the binding kinetics in greater detail using bio-layer interferometry (BLI), which measures the effective mass change at the surface of a sensor tip. As expected, the S2 protein control yielded no binding (Figure 1C, D). However, BLI tips loaded with RBD measured a dissociation constant (K_D_) of 750 pM for saRBD1, while tips coated with S1 yielded a K_D_ 1880 pM. S trimer loaded tips showed the strongest binding with a K_D_ of 674 pM, consistent with our ELISA results. These K_D_’s are lower than the previously reported 15 nM K_D_ of the RBD-ACE2 interaction, suggesting that saRBD-1 binds SARS-CoV-2 with at least an order of magnitude greater affinity than ACE2 (Glasgow et al., 2020).

**Figure 2:**
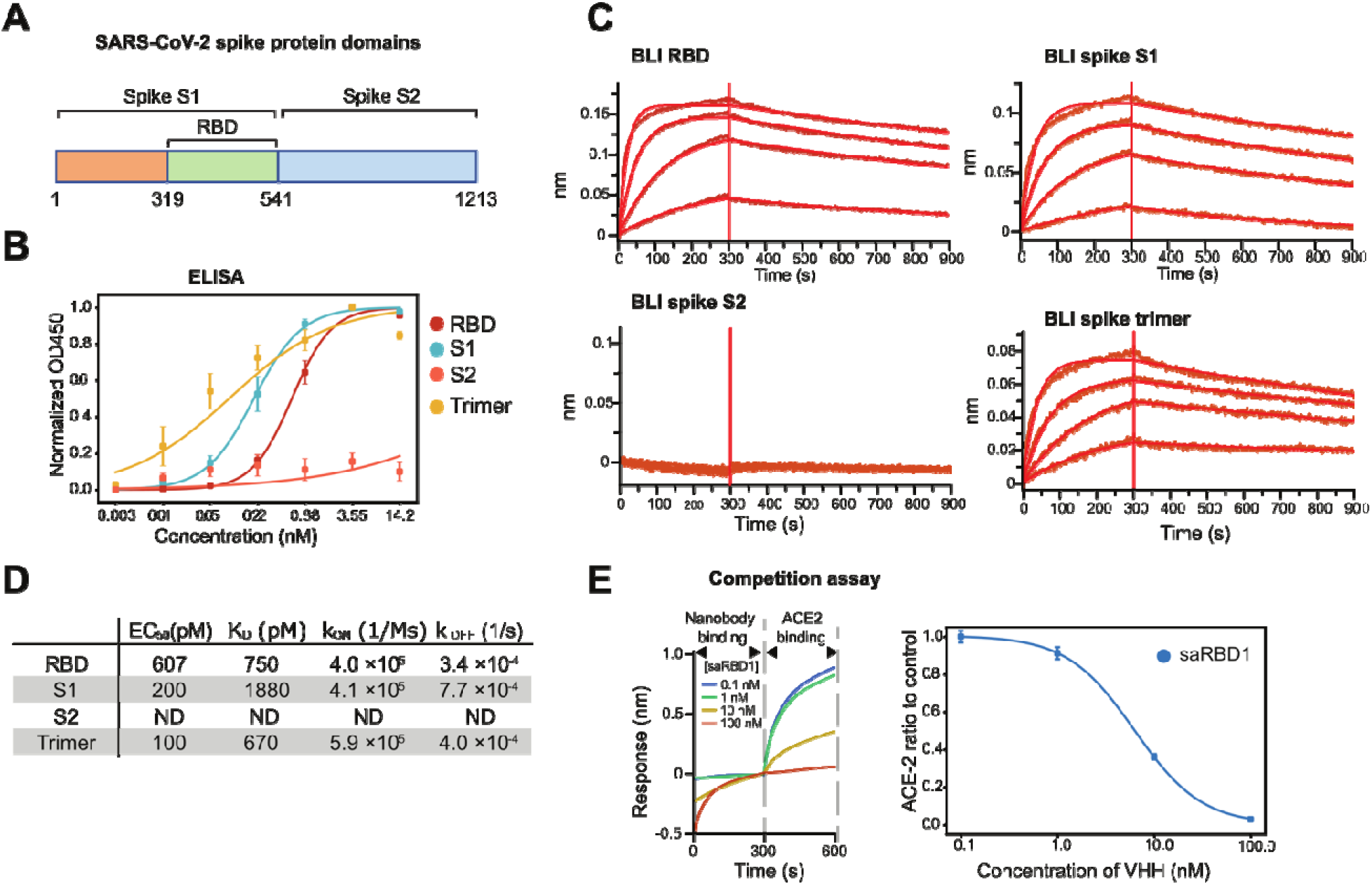
VHH saRBD-1 binds SARS-CoV-2 spike and receptor domain with high affinity. A) Schematic identifying SARS-CoV-2 S protein domains. B) ELISA binding assay of saRBD-1 on plates coated with SARS-CoV-2 RBD, S1, S2, and full-length S trimer where saRBD-1 is seen to bind RBD, S1, and full-length S trimer, but not S2. Curves show the average of 3 replicate experiments. C) Representative BLI curves of saRBD-1 binding kinetics experiments on SARS-CoV-2 S RBD, S1, S2, and full-length trimer where saRBD-1 is seen to bind RBD, S1, and full-length S trimer, but not S2. Biotinylated spike constructs were pre-bound to streptavidin biosensor tips, after which association and dissociation steps were carried out in saRBD-1 solutions at (from top to bottom): 100nM, 31.6nM, 10nM, and 3.16nM. D) Summary data of ELISA (B) and BLI (C) results. ELISA EC_50_ values and BLI K_D_, k_ON_ and k_OFF_ values are the average of at least two replicates. E) saRBD-1 competes with ACE2 receptor for binding SARS-CoV-2 S RBD. ACE2 binding to RBD after blocking with different concentrations of saRBD-1 by BLI. Resulting ACE2 binding values were by dividing by ACE2-only control. Left is representative BLI trace and right is normalized ACE2 binding values fit to a dose-response curve, average of two replicates. Error bars in all plots represent standard error.

To more thoroughly examine this, we performed BLI-based competition assays to determine if saRBD-1 is able to block the RBD-ACE2 interaction (Figure 2E). In this assay, BLI tips were first loaded with RBD followed by varying concentrations of saRBD-1 VHH to block the RBD binding sites before finally transferring to a solution with a fixed concentration of ACE2. We found that saRBD-1 bound competitively with ACE2, and that a concentration of 6 nM of saRBD-1 was sufficient to block 50% of ACE2 binding. These results indicated that saRBD-1 binds specifically to the RBD subunit of native trimeric S protein with picomolar affinity and blocks the subsequent interaction of RBD with ACE2.

### VHH saRBD-1 neutralizes both SARS-CoV-2 spike-pseudotyped lentiviruses and live SARS-CoV-2 virions

As an initial test of neutralization, we performed immunofluorescence microscopy on Vero E6 cells infected in the presence of 179 nM of either saRBD-1, which we found to completely block infection, or 179 nM control VHH52 (Figure 3A). Infection was visualized with the anti-dsRNA antibody. To quantify the neutralizing potency of saRBD-1 we performed experiments for both pseudotyped virus and live SARS-CoV-2. For pseudotyped virus neutralization assays, GFP-bearing SARS-CoV-2 S pseudotyped lentivirus was incubated with dilutions of saRBD-1 or control VHH52 before being added to target 293T-ACE2 cells. Successful lentivirus transduction was detected by high-content fluorescence microscopy of GFP signals. The 50% inhibitory concentration (IC_50_) of the VHH for this lentivirus challenge was 4.26 nM (Figure 3B) while VHH52 showed no inhibition. To control for non-specific inhibition of lentivirus transduction, lentivirus was generated pseudotyped with the VSV G protein in lieu of SARS-CoV-2 S; VSV G pseudotyped virus was not neutralized by saRBD-1 (Figure 3B).

**Figure 3:**
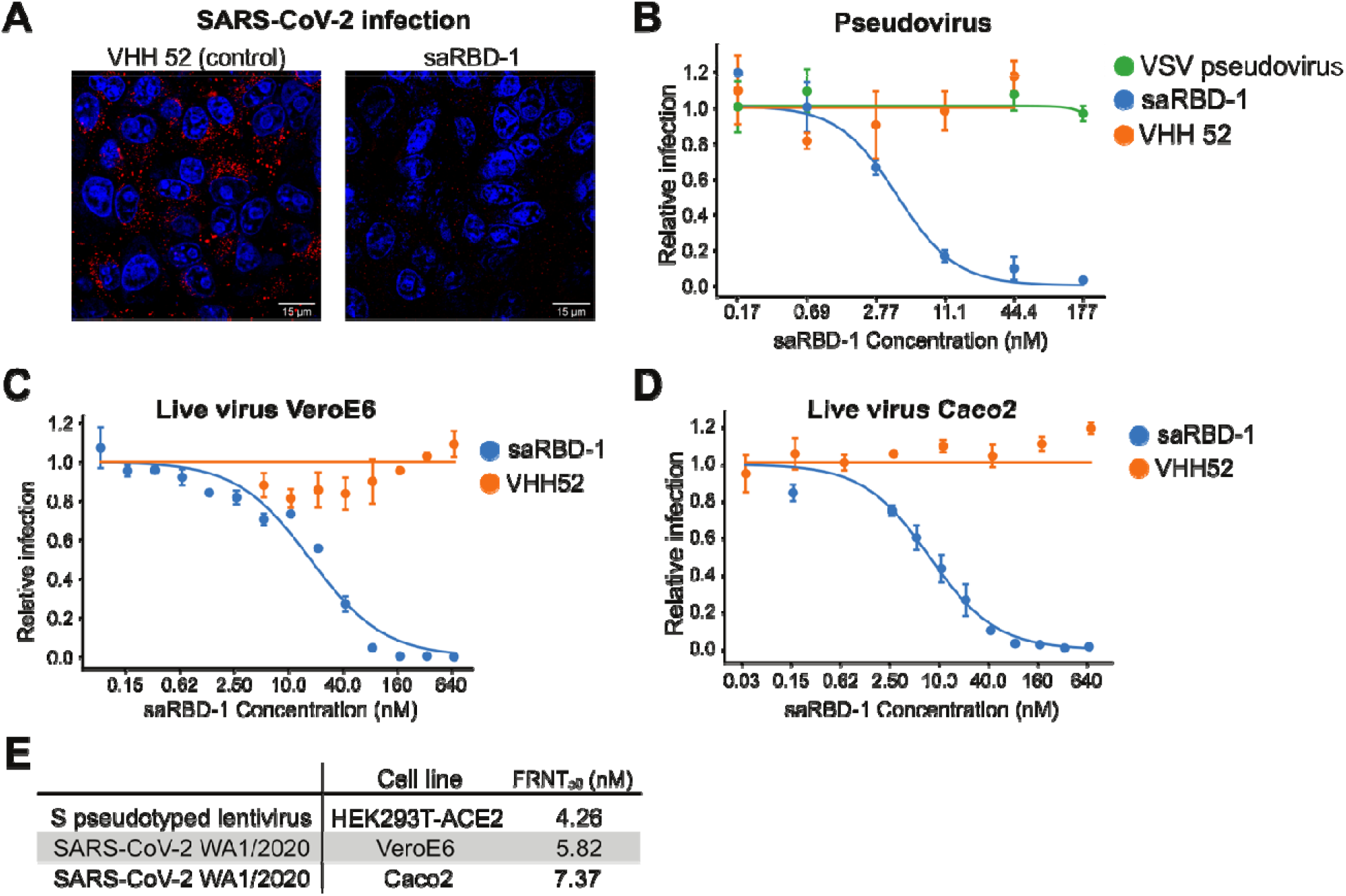
VHH saRBD-1 neutralizes both SARS-CoV-2 spike-pseudotyped lentiviruses and live SARS-CoV-2 virions. A) Representative images of assays used to quantify the effects of saRBD-1 on viral entry. Representative microscopy of SARS-CoV-2 dsRNA (red) in presence of 179 nM saRBD-1 or control VHH 52, cell nuclei stained with DAPI (blue). B) Neutralization of S-pseudotyped lentivirus by saRBD-1. ACE2 positive HEK-293T cells were infected with GFP reporter pseudovirus and either saRBD-1 or control VHH52. VSV G protein pseudovirus was incubated with saRBD-1 similarly to S-pseudovirus. Cells were fixed after 48 hours, then stained with DAPI and imaged. GFP signals were normalized to virus-only control wells. Averages of three replicate experiments are shown. Neutralization of live SARS-CoV-2 virus by saRBD-1 during infections of C) VeroE6 cells and D) Caco-2 cells. Neutralization was measured by focus forming assay of live WA1/2020 pre-incubated with saRBD-1 or control VHH52. Data represent the average of at least two replicate experiments, each in technical triplicate. E) Summary table of 50% focus reduction neutralization (FRNT_50_) results from pseudovirus and live virus neutralization assays. Error bars in all plots represent standard error.

To determine VHH inhibitory activities against live SARS-CoV-2 virus, focus forming assays were performed using SARS-CoV-2 WA1/2020 strain and saRBD-1. For the assay, Vero E6 or human colorectal epithelial (Caco-2) cells were infected with SARS-CoV-2, then stained with anti-S alpaca polyclonal sera as a primary antibody and an HRP-conjugated secondary antibody, facilitating visualization of SARS-CoV-2 infected cells (Figure 3C, D). The 50% focus reduction neutralization titer (FRNT_50_) was found to be 5.82 nM for Vero E6, and 7.4 nM for Caco2 (Figure 3E). In comparison, the non-neutralizing VHH52 failed to decrease foci. Thus, it is evident that monovalent saRBD-1 is a potent neutralizer of live SAR-COV-2 *in vitro,* even at low nanomolar concentrations.

### An Fc conjugated bivalent VHH construct, and a dimeric saRBD-1 construct show improved binding and neutralization of SARS-CoV-2

While monomeric saRBD-1 demonstrated exceptional neutralization of SARS-CoV-2, multimeric VHHs previously have been shown to have improved affinities and neutralization capabilities (Günaydın et al., 2016; Hanke et al., 2020; Schoof et al., 2020). To test this with saRBD-1, we utilized a mammalian vector to express saRBD-1 conjugated to human IgG Fc with a short hinge(Hanke et al., 2020; Tiller et al., 2008). The resulting chimeric protein is secreted as a dimer due to disulfide bridging of two Fc regions, and thus acts as a partially humanized heavy-chain only antibody (Figure 4A). This approach allows for improved binding due to avidity effects and greater steric blockage of the ACE2 binding site of the S protein. Simultaneously, we produced a bivalent construct of saRBD-1 (BI-saRBD-1) attached by a flexible (GGGGS)_4_ linker (Shan et al., 1999; Wrapp et al., 2020a). To determine binding kinetics of the saRBD-1 Fc-dimer (Fc-saRBD-1) to RBD, we utilized ELISA and BLI (Figure 4B-C, Figure S2). The EC_50_ of Fc-saRBD-1 as measured by ELISA was 392 pM, a 50% stronger affinity as compared to monovalent saRBD-1. The K_D_ of Fc-saRBD-1 as measured by BLI was 302 pM, primarily driven by a 3-fold reduction in the K_OFF_ compared to monovalent saRBD-1. Using our pseudovirus neutralization assay, the neutralization ability of the Fc-saRBD-1 dimer improved to an IC_50_ of 100 pM, over a 40-fold improvement compared to monomeric saRBD-1 (Figure 4D, F). Neutralization of live SARS-CoV-2 by Fc-saRBD-1 had an FRNT_50_ of 118 pM in VeroE6 cells and 218 pM in Caco2 cells, Bi-saRBD-1 had an FRNT_50_ of 243 pM in VeroE6 cells and 728 pM in Caco2 cells. Compared to monomeric saRBD-1, this represents a 49-fold (Fc-saRBD-1) and 24-fold (Bi-saRBD-1) improvement in neutralization on VeroE6 cells, and a 34-fold (Fc-saRBD-1) and 10-fold (Bi-saRBD-1) improvement in Caco2 cells (Figure 4E, F). The slightly improved neutralization shown by the Fc construct relative to the plain bivalent construct may be explained by the increased stearic hindrance from the bulky Fc portion (Hanke et al., 2020).

**Figure 4:**
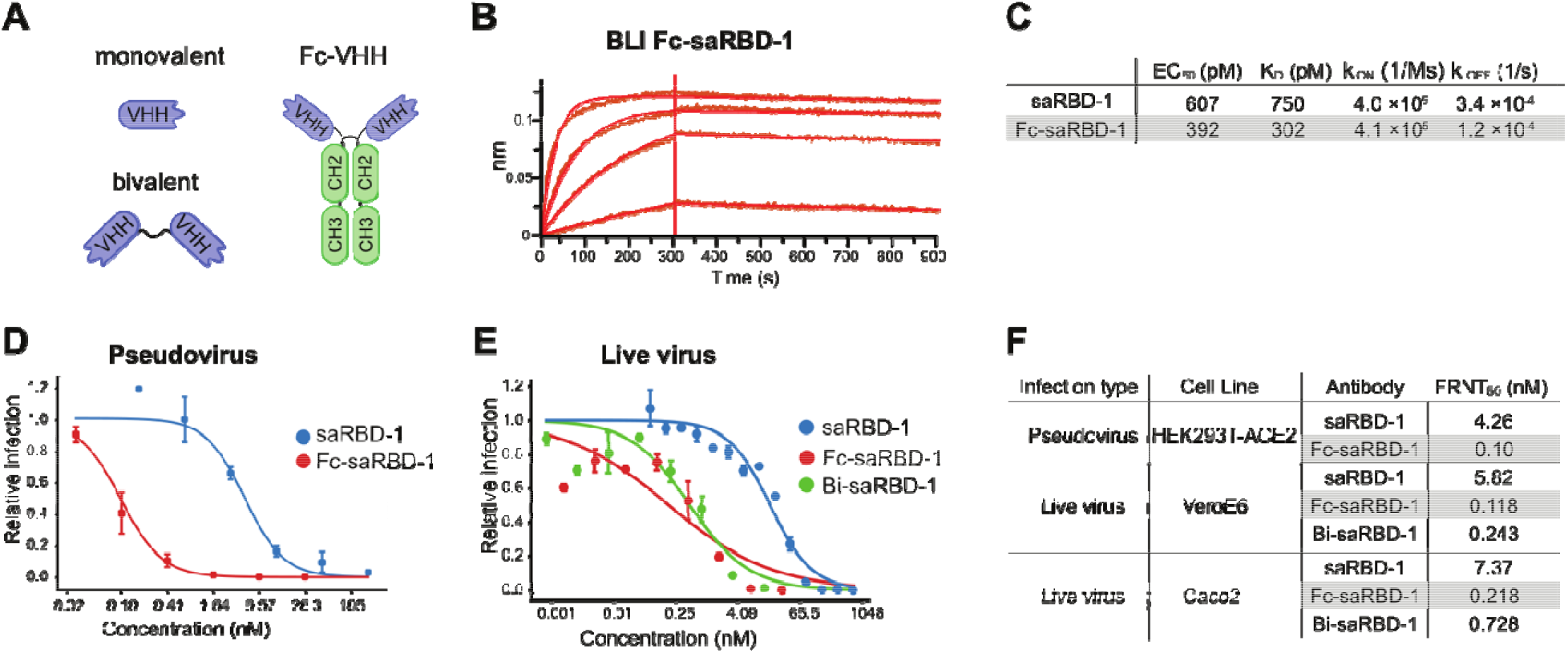
An Fc conjugated bivalent VHH construct, and a dimeric saRBD-1 construct show improved binding and neutralization of SARS-CoV-2. A) Schematic of monovalent, Fc-conjugated dimeric, and bivalent constructs. B) Representative BLI curves for Fc-saRBD-1 kinetic binding experiments on SARS-CoV-2 RBD. Biotinylated RBD was pre-bound to streptavidin biosensor tips, after which association and dissociation steps were carried out in saRBD-1 solutions at (from top to bottom): 100nM, 31.6nM, 10nM, and 3.16nM. C) Summary table of BLI kinetic parameters. Data are the average of two replicates. D) SARS-CoV-2 S pseudovirus neutralization curves showing the average of three microscopy experiments. E) Live SARS-CoV-2 (WA1/2020) neutralization curves showing the average of at least (n=2) replicate focus forming assay experiments, each in technical triplicate. F) Summary table of FRNT_50_ results from pseudovirus and live virus neutralization assays. Error bars in all plots represent standard error.

### saRBD-1 VHH is stable and maintains its activity after heat treatment, lyophilization and nebulization

One of the major advantages of VHHs over conventional antibodies is their inherent stability. We evaluated the stability of saRBD-1 by subjecting it to some of the conditions that are likely to be encountered during production, transport, and delivery of protein-based therapeutics we evaluated the stability of saRBD-1 in elevated temperature, lyophilization, and nebulization. We treated VHH to each condition, then measured of protein loss, binding kinetics, and neutralizing ability of the treated VHH aliquots. Aliquots of saRBD-1 were incubated for 1 hour at 50°C then centrifuged to remove aggregates before measurement of protein loss by OD_280_, which showed a 19% reduction. The treated aliquots were then checked by BLI on RBD (Figure 5G, I), which showed minimal loss of activity concomitant with the reduction in measured protein concentration. Similar measurements were performed using lyophilized (29% protein loss) and nebulized (77% protein loss) samples. Nebulization is known to be a harsh process, particularly when performed in unmodified PBS solution with a jet nebulizer, and our numbers mirror previous reports of 4-fold loss of activity after nebulization with an ultrasonic nebulizer (Schoof et al., 2020). In total, we found that the K_D_ was 938 pM for heat treatment, 936 pM for lyophilized, and 3.65 nM for aerosolized, amounting to 1.25-fold, 1.25-fold, and 4.8-fold increases respectively, which align with our protein loss determinations.

**Figure 5:**
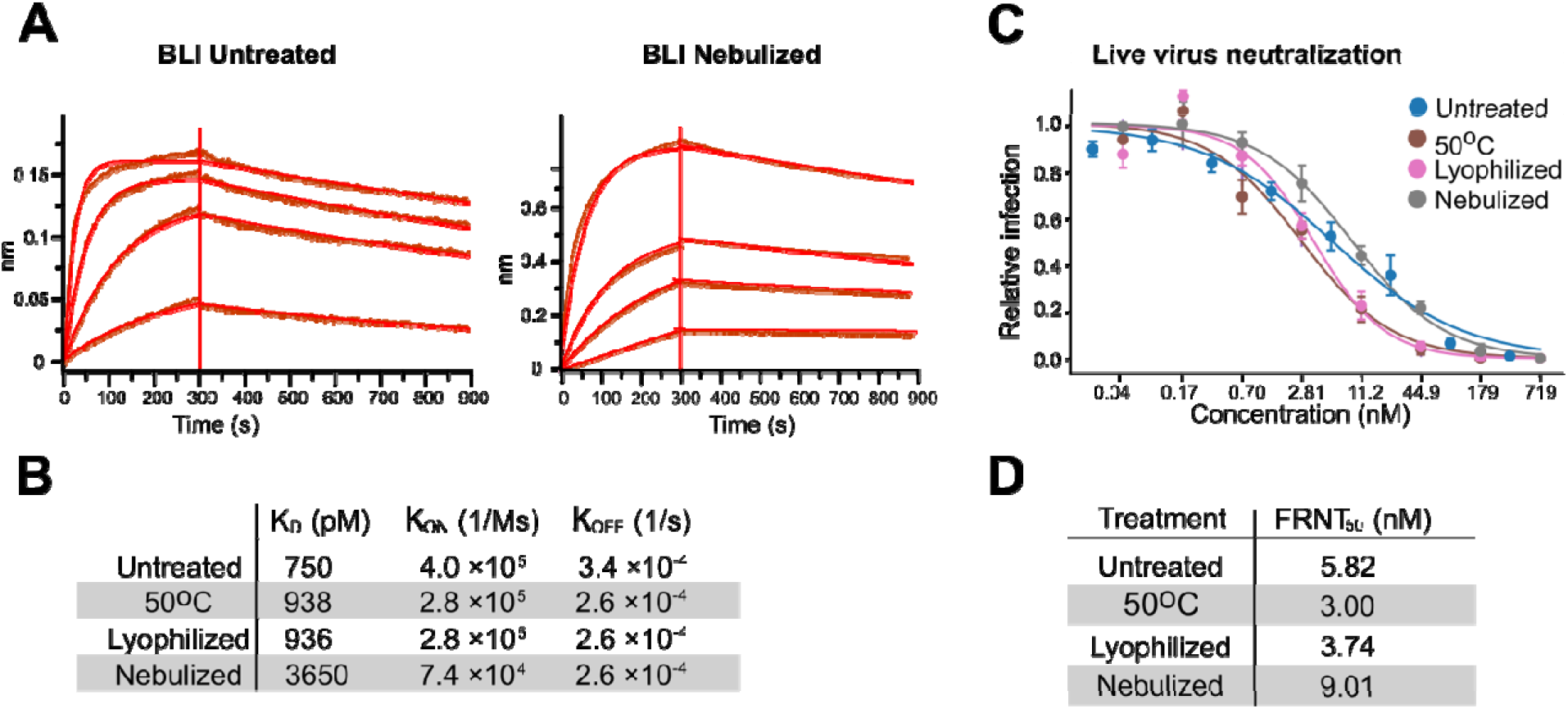
saRBD-1 VHH is stable and maintains its activity after heat treatment, lyophilization and nebulization. A) Representative BLI curves of kinetics experiments of saRBD-1 binding RBD of untreated and nebulized samples. Biotinylated RBD was pre-bound to streptavidin biosensor tips, after which association and dissociation steps were carried out in saRBD-1 solutions at (from top to bottom): 100nM, 31.6nM, 10nM, and 3.16nM. B) Summary table of BLI kinetics experiments of untreated, heat treated, lyophilized, and nebulized saRBD-1 samples. Data are the average of two replicates. C) Live SARS-CoV-2 focus forming assay neutralization curves for untreated, heat treated, lyophilized, and nebulized saRBD-1 samples, showing the average of at least two replicate experiments, each in technical triplicate. E) Summary table of FRNT_50_ results from live virus neutralization assays. Error bars in all plots represent standard error.

To assay effects of these treatments on neutralizing activity, we carried out focus forming assays in VeroE6 cells utilizing the heat treated, lyophilized, and nebulized saRBD-1 samples (Figure 3H, J). We found that 50°C treated, lyophilized, and nebulized saRBD-1 yielded FRNT_50_s of 3.00 nM, 3.74 nM, and 9.01 nM respectively. In comparison, untreated saRBD-1 yielded a FRNT_50_ of 5.82 nM. Therefore, only nebulization reduced saRBD-1 neutralizing capability, with a 1.56-fold reduction. Overall saRBD-1 appears functionally stable, and it maintains nanomolar neutralization activity towards RBD even after destabilizing treatments.

### SaRBD-1 effectively neutralizes SARS-CoV-2 variants of concern

Because of the prevalence of SARS-CoV-2 variant strains of concern (VOCs) significantly divergent from the base strain (Figure 6A), we sought to test saRBD-1’s affinity for mutated RBD-N501Y and neutralizing abilities against clinical VOC isolates We generated a variant RBD to test saRBD-1-RBD interactions. Using site directed mutagenesis, we created a spike and RBD variant that contained the N501Y mutation found in several of the circulating VOCs (Figure 6B). Using BLI, we found binding of saRBD-1 to RBD-N501Y was similar to WT saRBD-1 (Figure 6C, D), with a K_D_ of 767 pM compared to the WT value of 750 pM (Figure 6E). The affinity of saRBD-1 against both wild-type and mutant RBD constructs were stronger than the 15 nM affinity of RBD for ACE2 (Glasgow et al., 2020), indicating that the N501Y amino acid change is unlikely to affect neutralization.

**Figure 6:**
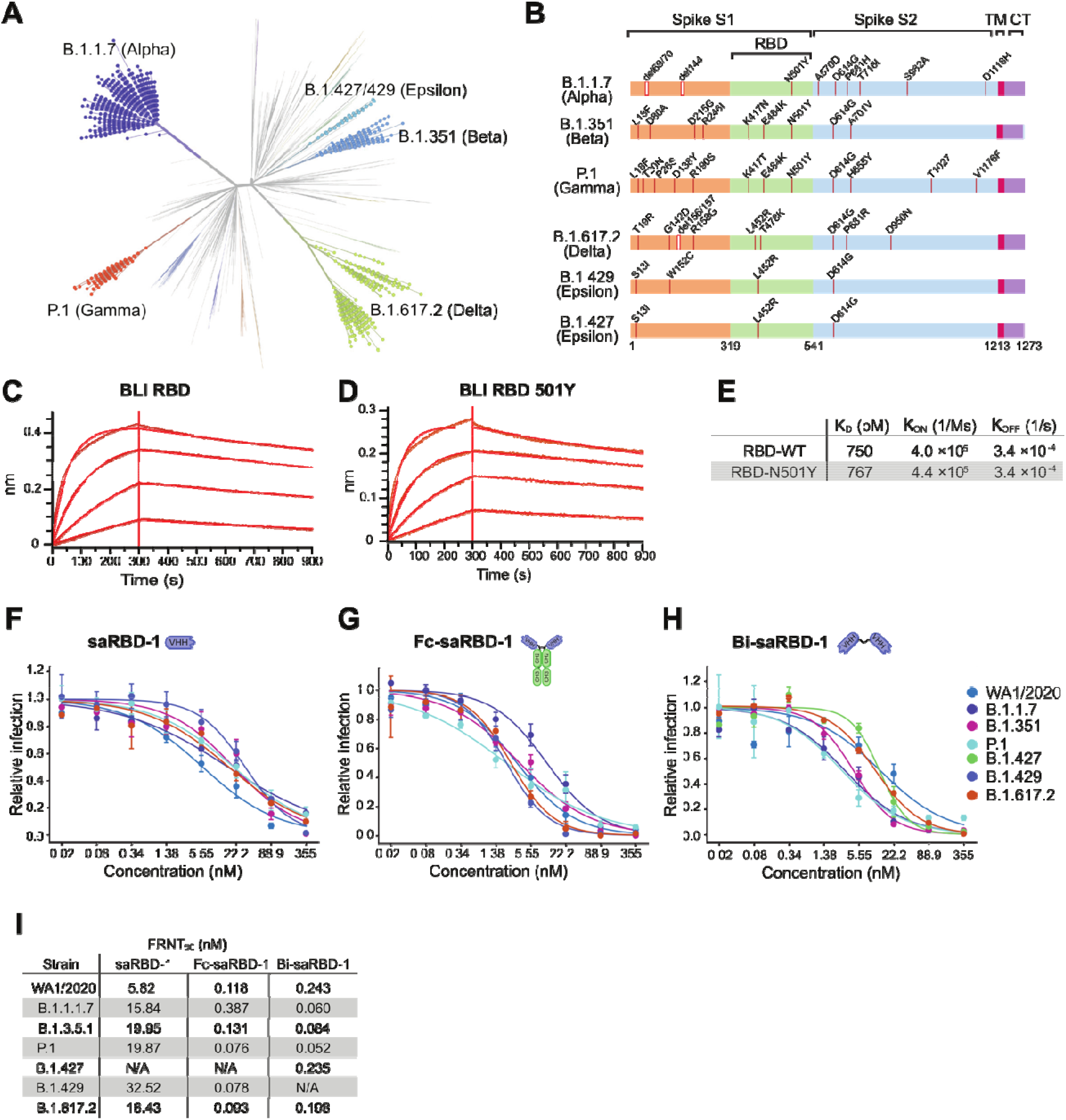
SaRBD-1 effectively neutralizes SARS-CoV-2 variants of concern. A) Phylogenetic tree including the VOCs. The tree was generated in Nextstrain of all available variants with the VOCs used in this study highlighted and labeled. B) Schematic diagram of the variant spike protein amino acid changes present in the VOCs. Representative BLI curves of kinetic experiments of saRBD-1 on C) RBD and D) RBD-N501Y. Biotinylated RBD and RBD-N501Y were pre-bound to streptavidin biosensor tips, after which association and dissociation steps were carried out in saRBD-1 solutions at (from top to bottom): 100nM, 31.6nM, 10nM, and 3.16nM. E) Summary table of binding kinetic values of saRBD-1 on RBD and RBD-N501Y, determined by BLI. Data are the average of two replicates. Live SARS-CoV-2 focus forming assay neutralization curves of the VOCs for F) monomeric saRBD-1, G) dimeric Fc-saRBD-1, and H) bivalent Bi-saRBD-1. Data are the average of two replicate experiments, each in technical triplicate. I) Summary table of FRNT_50_ results from live virus neutralization assays. N/A: Not tested. Error bars in all plots represent standard error.

To more directly address variant cross-neutralization, we obtained clinical isolates of the following SARS-CoV-2 VOCs with known RBD mutations: Alpha containing N501Y; Beta containing K417N, E484K, N501Y; Gamma containing K417T, E484K, N501Y; Epsilon containing L452R; and Delta containing RBD L452R and T478K. Compared to the FRNT_50_ of 5.82 nM on WA1/2020, the variants were neutralized with FRNT_50_ values of 15.84 nM (Alpha), 19.95 nM (Beta), 19.87 nM (Gamma), 32.52 nM (Epsilon), and 16.43 nM (Delta), representing 3-fold to 6-fold reductions that may be due to marginal differences in binding or natural experimental variation in the focus forming assays (Figure 6F-I). We additionally sought to test the efficacy of our Fc-conjugated VHH against all VOCs. We found FRNT_50_’s of 118 pM (WA1/2020), 387 pM (Alpha), 131 pM (Beta), 76 pM (Gamma), 78 pM (Epsilon), and 93 pM (Delta), which are all within 3-fold of WA1/2020 (Figure 6I). Finally, we utilized our Bi-saRBD-1 construct for VOC neutralization assays. With this construct, we found FRNT_50_’s of 243 pM (WA1/2020), 60 pM (Alpha), 56 pM (Beta), 235 pM (Gamma), 198 pM (Delta), where all variants are better neutralized than WA1/2020. Overall, our monomer saRBD-1 displayed low nanomolar FRNT_50_’s against all VOCs, while our dimeric constructs retained picomolar levels. Therefore, saRBD-1 likely targets an RBD epitope that is conserved across all tested SARS-CoV-2 VOCs.

### SaRBD-1 competes with class-1 monoclonal antibody B38

To narrow down the binding epitope of saRBD-1, we performed competitive binding assays against monoclonal antibodies from the three primary classes of RBD binding antibodies (Figure 7A). Class 1 antibodies bind epitopes around K417 and tend to bind spike in the up conformation, while class 2 antibodies bind epitopes around E484 and can bind both up and down conformations, and class 3 antibodies bind epitopes around L452, distal to the ACE2 contact surface (Barnes et al., 2020). We selected representative antibodies from each class: class 1: B38 (Wu et al., 2020), class 2: Ly-Cov555 (Greaney et al., 2021; Jones et al., 2021), class 3: REGN10987 (Greaney et al., 2021; Weinreich et al., 2021) to use in a biolayer interferometry (BLI) competitive binding assay. In this assay, SARS-CoV-2 spike RBD protein was attached to a sensor and first exposed to saRBD-1, which bound strongly during the first 300 seconds. In the subsequent step, the sensors were transferred to solutions containing the representative monoclonal antibodies. SaRBD-1 successfully blocked B38 from binding, indicating that they likely bind to overlapping epitopes (Figure 7B). The class 2 & 3 antibodies were not affected by saRBD-1. A control experiment confirmed that B38 binds successfully when saRBD-1 was absent (Figure 7C). These results were recapitulated with a dimeric Fc-saRBD-1 construct (Figure7D). Hence, saRBD-1 is most likely a class 1 binder. An unlikely alternative is that saRBD-1 binds a distal site non-competitive with the class 3 antibody, but forces RBD into a down conformation unsuitable for B38 binding.

**Figure 7:**
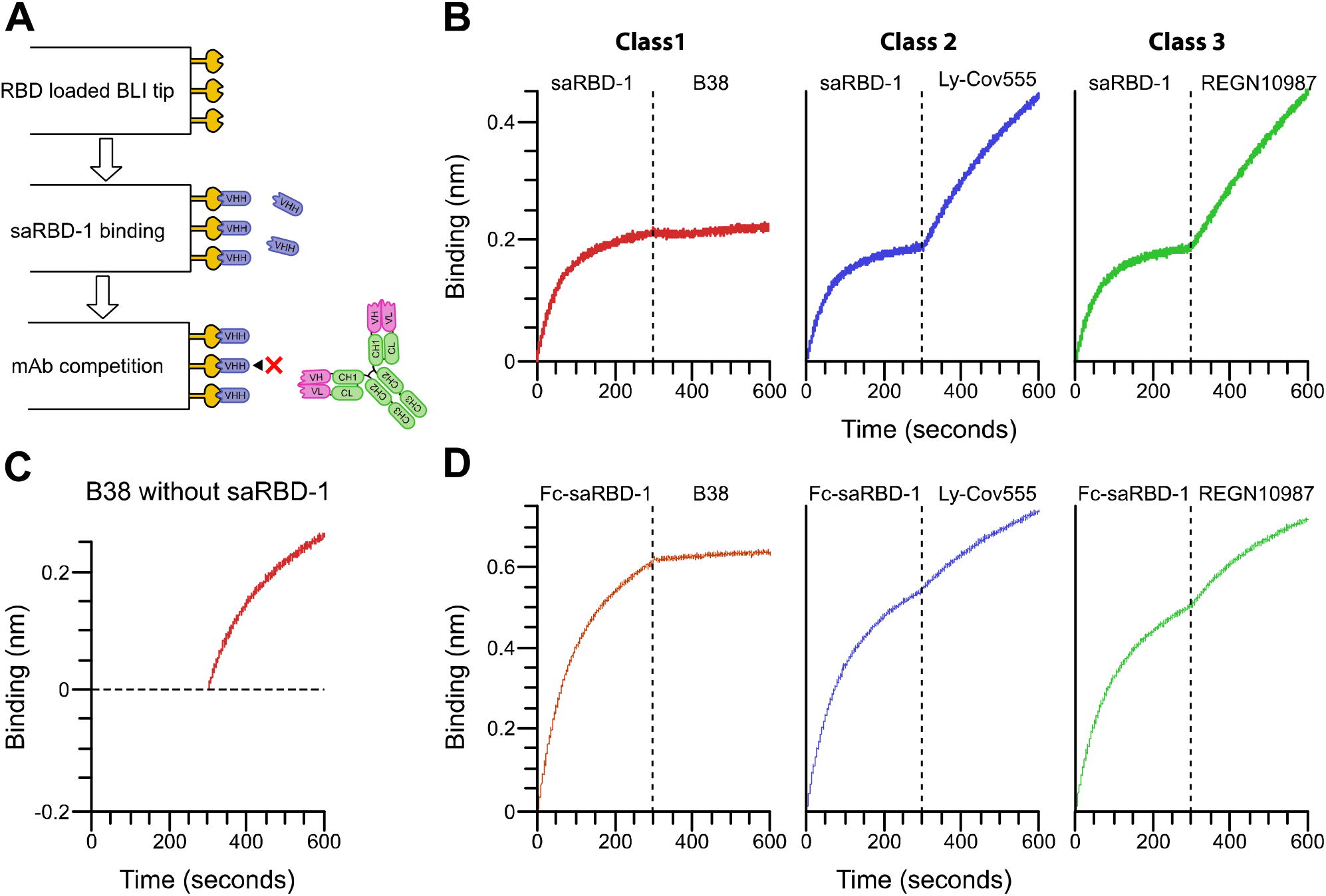
saRBD-1 competes with class-1 monoclonal antibody B38. A) Experimental design of BLI-based competition assay of monoclonal antibodies. B) BLI measurement of saRBD-1 binding followed by class 1, 2, or 3 monoclonal antibodies. Binding. B38 (class 1) is shown in red, LyCoV-555 (class 2) is in blue, and REGN19087 (class 3) is in green. C) BLI measurement of B38 (class 1) binding in the absence of saRBD-1. D) BLI measurement of Fc-saRBD-1 binding followed by class 1, 2, or 3 monoclonal antibodies.

## Discussion

The ongoing SARS-CoV-2 pandemic is a threat to global public health; the discovery and synthesis of additional therapeutics and vaccines are needed to address this. Although the scientific community has developed effective vaccines (Baden et al., 2021; Daniel et al., 2021; Shen et al., 2021) against the initial SARS-CoV-2 outbreak, ongoing concern that a vaccine resistant VOC could result in a resurgent outbreak has played out with the arrival of the Omicron VOC (Liu et al., 2021c; Zhang et al., 2021). Two of the most widely utilized vaccine options from Pfizer/BioNTech and Moderna, are expensive and unstable mRNA vaccines, requiring specialized transportation and storage (Crommelin et al., 2021; Kartoglu et al., 2020). This has resulted in a dearth in vaccine availability for communities around the world compounding the human impact of the SARS-CoV-2 pandemic and transmission of VOCs (Holder; Mathieu et al., 2021). Increasing evidence also points to waning immune responses to the vaccines, increasing the risk of breakthrough infections (Levin et al., 2021; Shrotri et al., 2021) As such, novel therapeutics and vaccines should fulfill both the following conditions: 1) be affordable to produce, transport and store. 2) provide highly effective long-term protection against circulating VOCs. Our saRBD-1 VHH is an ideal match due to its cheap manufacture of bacterial purification, thermostability, and efficacy at VOC neutralization.

The ability of saRBD-1 to potently neutralize SARS-CoV-2 is critical to its potential. Antiviral VHHs have utility as prophylactics or therapeutics against viral infections (Ingram et al., 2018; Laursen et al., 2018). Hence, strongly neutralizing VHHs against SARS-CoV-2 are desirable. Other groups have isolated VHH candidates that bind S RBD and neutralize SARS-CoV-2 infections in situ (Güttler et al., 2021; Hanke et al., 2020, 2022, 2022; Koenig et al., 2021; Schoof et al., 2020; Wagner et al., 2021; Wrapp et al., 2020a; Xiang et al., 2020; Xu et al., 2021) and in animal models (Kim et al., 2021; Pymm et al., 2021; Wagner et al., 2021). The monomeric form of saRBD-1 potently neutralizes ancestral SARS-CoV-2 and VOCs with FRNT_50_ of around 5.82 nM. Other VHHs within a similar range of neutralizing potency have been reported (Pymm et al., 2021; Xu et al., 2021). Strong inhibition of SARS-CoV-2 is critical for VHH therapeutic potential, as the highest possible neutralizing strength is ideal for minimizing the effective dose of a potential treatment. Another encouraging quality of saRBD-1 is its extreme stability against multiple forms of insult. SaRBD-1 retained neutralizing activity and RBD-binding capability when heated to 50°C, nebulized or lyophilized. These tests are relevant because they mimic the likely transport, storage, and delivery conditions that are likely to be encountered by a therapeutic anti-SARS-CoV-2 nanobody.

As a bivalent construct or when conjugated to a human IgG Fc domain, saRBD-1 has comparable neutralizing capabilities to highly neutralizing monoclonal antibodies, reaching ∼100 pM FRNT50s against live SARS-CoV-2. As such, Bi-saRBD-1 and Fc-saRBD-1 may prove useful for prophylaxis in a similar manner to convalescent plasma transfusions, which have had positive clinical outcomes throughout the COVID-19 pandemic (Hu et al., 2020; Zeng et al., 2020). Although the addition of the Fc domain to nanobodies undermines the key beneficial features of small size and may cause antibody-dependent enhancement (Eroshenko et al., 2020), Fc-conjugation is known to significantly increase the in vivo half-life of nanobodies (Rotman et al., 2015).

The most appealing characteristic of saRBD-1 is its activity against SARS-CoV-2 VOCs. VOCs pose a global threat beyond that of the initial SARS-CoV-2 pandemic, as such effective treatments are especially valuable to protecting public health. VOCs tend to have increased transmission compared to the initial pandemic strains of SARS-CoV-2 (Chen et al., 2021; Korber et al., 2020; Liu et al., 2021a). Furthermore, VOCs with multiple S protein RBD mutations resist neutralization by vaccinated sera and convalescent patient sera (Bates et al., 2022; Chen et al., 2021; Liu et al., 2021c). Development of VOCs increases the possibility of new mutations developing that escape vaccination, especially with partial vaccination of global population or waning antibody levels in those vaccinated (Levin et al., 2021; Mathieu et al., 2021; Shrotri et al., 2021). Although several VHHs are reported with neutralizing activities against Alpha (Pymm et al., 2021; Zupancic et al., 2021) and Beta (Güttler et al., 2021; Hanke et al., 2022; Mast et al., 2021; Wagner et al., 2021; Xu et al., 2021; Zupancic et al., 2021), few are published with activity against Gamma (Mast et al., 2021) and Delta (Wagner et al., 2021). Previous studies have reported that combinations of neutralizing VHHs delivered *in-situ* SARS-CoV-2 infections are important to suppress development of escape mutations (Wrapp et al., 2020a) and better neutralize variant strains (Pymm et al., 2021). A great variety of VOC neutralizing VHHs will be crucial if nanobodies are to play a role as SARS-CoV-2 therapeutics against future evasive VOCs similar to Omicron. Thus saRBD-1 can be valuable due to its proven ability to neutralize 5 distinct VOC strains, including Delta.

Neutralizing antibodies against SARS-CoV-2 RBD can be organized into three classes depending on their targets in the RBD-ACE2 interface and confirmation of RBD when binding. Class 1 binds up RBD at the ACE2 binding site, class 2 binds up and down RBD at the ACE2 binding site, and class 3 binds up and down through residues distal to the ACE2-binding site (Barnes et al., 2020). Therefore, class 1 and 2 antibodies target RBD residues that are frequently mutated in VOCs such as K417, E484 and N501; L452 is the most commonly mutated residue bound by class 3 antibodies (Greaney et al., 2021). Among the published VHHs, Fu2 (Hanke et al., 2022) and WNb2 (Pymm et al., 2021) appear to be class 1, while Nb6 and Nb11 (Schoof et al., 2020), Nb20 (Xiang et al., 2020) and Ty1 (Hanke et al., 2020) are class 2. Class 3 nanobodies are also reported, including WNb10 and 15 (Pymm et al., 2021), Nb12 and Nb30 (Xu et al., 2021), and VHH72 (Wrapp et al., 2020a). We determined that saRBD-1 belongs to class 1 due to competition with class 1 monoclonal antibody B38 (Wu et al., 2020). Interestingly, saRBD-1 neutralizes VOCs containing K417, E484, and N501 mutations that typically affect class 1 and 2 antibodies, suggesting its epitope identity or mechanism of neutralization may be atypical for class 1 neutralizing antibodies. The recently published VHH Fu2 is an example of an atypical mechanism of SARS-CoV-2 neutralization (Hanke et al., 2022). Fu2 binds as a class 1 antibody to block ACE2 biding to RBD, yet simultaneously induces dimerization in full-length spike to further disrupt ACE2 interactions. Fu2 was also found to neutralizes the Beta variant without significant loss of potency. Thus, a VHH may have unexpected levels of utility outside of those predicted by epitope class, which can aid in binding mutated RBD variants.

We found that saRBD-1 binds competitively with human ACE2 for SARS-CoV-2 spike RBD, and that pre-incubation of RBD with saRBD1 blocks ACE2 binding, a necessary step to infection. Low nanomolar concentrations of monovalent saRBD-1 successfully neutralize clinical isolates of the Alpha, Beta, Gamma, Epsilon, and Delta VOC as a likely class 1 antibody. Both the Bi-saRBD-1 and Fc-saRBD-1 demonstrate improved binding, and they successfully neutralize the variants at picomolar concentrations with no discernable loss of potency. Due to its high neutralizing efficacy, saRBD-1, alone or in combination with other ultra-potent VHHs, is an excellent candidate for development into a therapeutic to manage severe COVID-19.

## Methods

### Key resource table

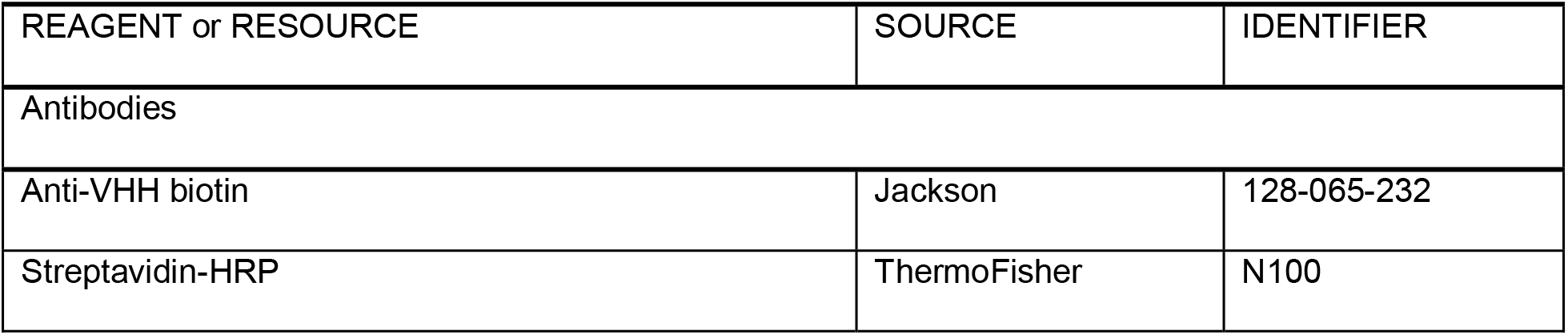
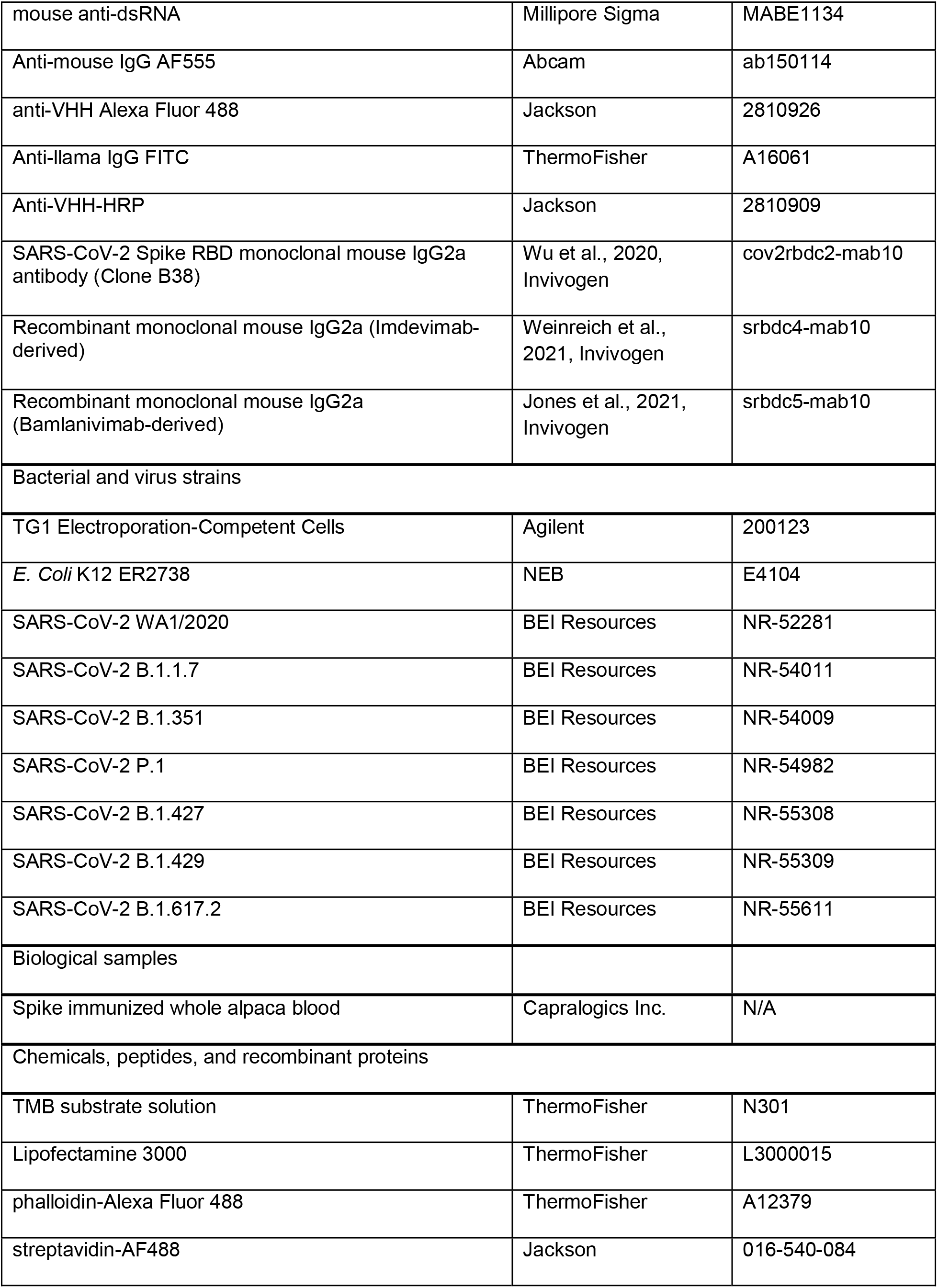
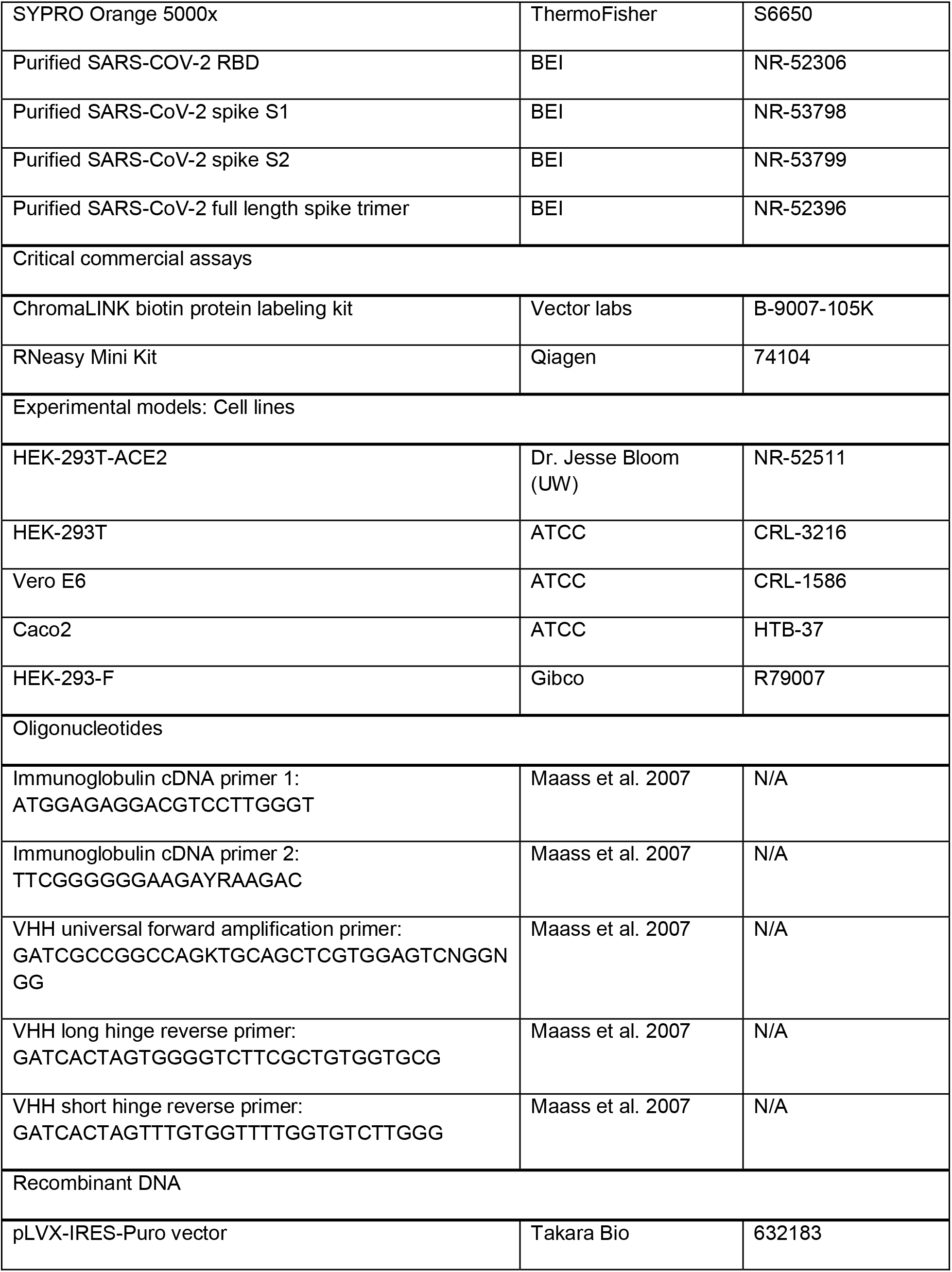
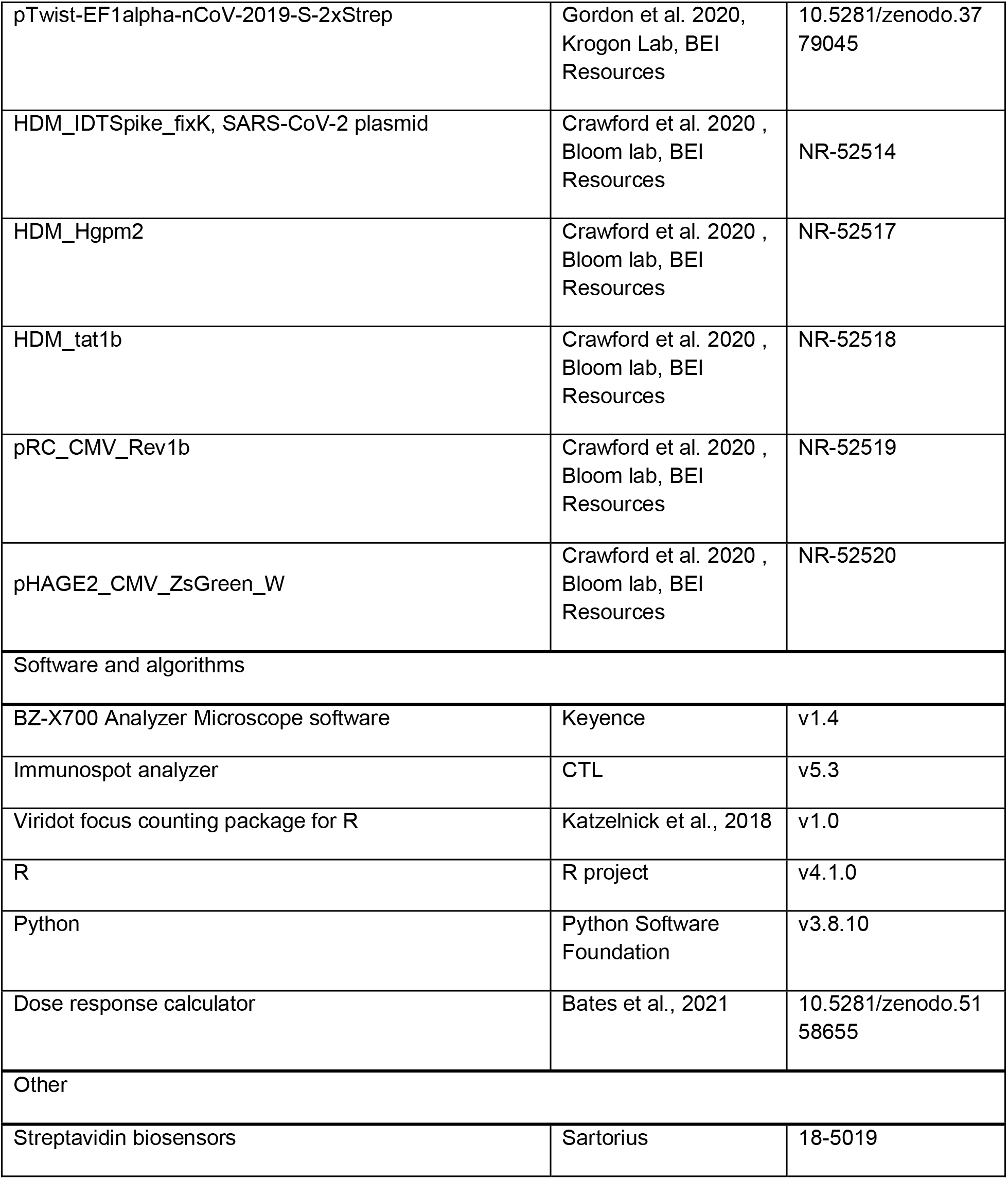

### Experimental model and subject details

HEK-293T stable cell lines expressing human ACE2 receptor (HEK-293T-ACE2) were a kind gift from Dr. Jesse D. Bloom from University of Washington, and described previously (Crawford et al., 2020). Low-passage HEK-293T, HEK-293T-ACE2, and Vero E6 cells were cultured in D10, which consisted of Dulbecco’s Modified Eagle Medium (DMEM) supplemented with 10% fetal bovine serum (FBS), 1% Penn-Strep, 1% non-essential amino acids (NEAA). 37°C. Caco2 cells were cultured in D20 (D10 with 20% FBS instead of 10%). Cells were cultured in T75 dishes, passaged with Trypsin at 95% confluency to avoid overcrowding.

### RBD protein purification and biotinylating

Purified SARS-CoV-2 S-RDB protein was prepared as described previously (Amanat et al., 2020). Briefly, codon optimized his-tagged RBD or RBD containing the N501Y mutation in pLVX-IRES-puro plasmid (Takare Bio) was used to make lentivirus vectors in HEK-293T cells, which were then used to infect HEK-293F suspension cells. The suspension cells were grown in FreeStyle^TM^ 293 expression medium (Gibco) for 3 days with shaking at 37°C with 8% CO_2_. Cell supernatants were collected, sterile filtered, and purified by Ni-NTA chromatography (Bierig et al., 2020). Purified protein was then buffer exchanged into phosphate-buffered saline (PBS) by dialysis and concentrated with 10kDa cutoff centrifuge filters (Millipore). For use in BLI, purified RBD was biotinylated using the ChromaLINK biotin protein labeling kit (Vector) according to the manufacturer’s instructions with 5x molar equivalents of labeling reagent to achieve 1.92 biotins/protein.

### VHH gene library construction

Alpacas were immunized at Capralogics Incorporate (Hardwick, MA). Animals received three immunizations of 1 mg purified SARS-CoV-2 RBD spaced three weeks apart. Blood was harvested 5 days after third immunization. Upon receipt PBMCs were isolated (Eppendorf) and used in an RNA extraction with Qiagen RNeasy mini kit (Qiagen). PBMC RNA, containing VHH genes, was converted into cDNA using Superscript III reverse transcriptase (Invitrogen). VHH genes were amplified with custom primers specific to short-hinge and long hinge VHH genes appended with Not1 and Asc1 restriction sites (Maass et al., 2007; Schmidt, 2018). The amplified gene mixture was cloned into a phage-mid plasmid derived from pCANTA5BE, then transformed via electroporation into bacteriophage competent TG1 Escherichia coli for production of a VHH displaying bacteriophage library. The pooled library was rescued at 37 C with shaking and plated. Plates containing serial dilutions were used to estimate total bacterial population, and therefore library diversity. A representative sample of 96 VHH clones were sent for sanger sequencing (Genewiz) to confirm VHH diversity.

### VHH panning

M13-derived helper phage were produced through standard protocols (Frei and Lai, 2016). VHH libraries in TG1 cells were transduced with helper phage to produce bacteriophage displaying individual VHHs. Phage were isolated through precipitation using 20% w/v polyethylene glycol 8000, then were resuspended in PBS. Phage libraries were then panned against full length stabilized SARS-CoV-2 spike protein trimer BEI resources NR-52396 at 10ug/ml alongside BSA controls. Bacteriophage were bound to the antigen, washed, then eluted with 200mM glycine pH 2.2, neutralized with 1M Tris pH 9.1, and transduced into ER2738 bacteriophage competent bacteria which were plated on antibiotic agar and incubated for 18 hours at 37 C. Panning success was determined by enrichment 10-fold greater total bacterial colonies above control panning background. Panning colonies were then pooled, and the protocol was repeated for 2^nd^ round panning using 2ug/ml antigen coating. A selection of colonies from 2^nd^ round panning were picked and grown up in 96 deep-well plates for screening.

### Screening of VHH candidates

96 well ELISAs coated with 1ug/ml purified RBD were used to determine RBD binding affinities of panning hits. ELISAs were ran with bacterial supernatant containing secreted VHHs as primary antibody, then anti-nanobody biotin antibody (Jackson #128-065-232) and streptavidin-HRP antibody (Thermo #N100) were used as secondary. 3,3’, 5’5”-tetramethylbenzidine (TMB) (ThermoFisher Scientific) was used as peroxidase substrate, 50 μl added for 10 minutes at room temperature (RT) then 50 μl of 2N H2SO4 was added as a stopping solution. Plate absorbance at 405 nm was measured using a CLARIOstar® Plus plate fluorimeter (BMG Labtech). VHH candidates with binding greater than 2-fold above average background were picked and sent for sanger sequencing (Genewiz) to identify VHH CDR3 regions. Clones that appeared multiple times in sequencing were cloned into a periplasmic expression vector (pHEN) by Gibson assembly. Bacteria were grown up in terrific broth (2% tryptone, 1% yeast extract, 90mM phosphate), induced with isopropylthio-β-galactoside (IPTG) at 30 C overnight. VHHs were isolated by osmotic shock (Saerens et al., 2004) , periplasmic fraction was isolated, and his-tagged VHHs were purified with Ni-NTA chromatography. Purified VHH was then buffer exchanged and concentrated with 3kDa cutoff centrifuge filters (Millipore), then filter sterilized by 22 μm centrifugal sterile filter (Millipore Sigma) prior to use in experiments.

### Multivalent saRBD-1 construction and purification

Fc-conjugated (Hanke et al., 2020) and bivalent saRBD-1 (Wrapp et al., 2020b) constructs were synthesized using guidance from prior publications. Fc-saRBD-1 gene was directly synthesized containing an sp6 promotor, secretion signal, saRBD-1, flexible hinge, human IgG1 Fc region (Tiller et al., 2008), and 6x his tags for cloning into pLVX-IRES-puro plasmid (Takare Bio). Lentivirus encoding Fc-saRBD-1 was produced, and protein was purified from transduced 293F cells as described for RBD purification above. Fc-saRBD-1 was then further purified by ion-exchange chromatography for a final yield of ∼4.6 mg per 100 ml of initial supernatant. Bivalent-saRBD-1 gene was synthesized containing two saRBD-1 genes separated by a flexible 20 a.a (GGGGS)_4_ linker, for cloning into pET24a bacterial expression plasmid. Bacteria were grown in terrific broth (2% tryptone, 1% yeast extract, 90mM phosphate), induced with isopropylthio-β-galactoside (IPTG) at 30°C overnight, lysed in Tris-NaCl buffer (500mM NaCl 20mM Tris, pH 8) by sonication and purified by Ni-NTA chromatography. Both multivalent proteins were buffer exchanged to remove excess imidazole and concentrated with 10 kDa cutoff centrifuge filters (Millipore), then filter sterilized by 22 μm centrifugal sterile filter (Millipore Sigma) prior to use in experiments.

### Cell transfection

HEK-293T cells seeded at 70-90% cell density, then transfected using Lipofectamine 3000 (ThermoFisher Scientific) as per manufacturer’s instructions. For S transfection, the SARS-CoV2 structural protein plasmid pTwist-EF1alpha-nCoV-2019-S-2xStrep a kind gift from the Krogan lab at UCSF, was used as described previously (Gordon et al., 2020). For pseudotyped lentivirus production, the following reporter plasmids and lentivirus packaging plasmids were used as described previously (Crawford et al., 2020): HDM_Hgpm2, HDM_tat1b, PRC_CMV_Rev1b packaging plasmids, SARS_CoV-2 S plasmid HDM_IDTSpike_fixK, and LzGreen GFP-reporter plasmid. Packaging, SARS-CoV-2 S, and reporter plasmids were a kind gift from the Bloom Lab at University of Washington. Per 6 well plate, 0.44 μg of each, packaging plasmid, 0.68 μg of S, and 2 μg of reporter plasmids were used for transfection. For all transfections, media was carefully removed 6 hours post transfection, and replaced with D10.

### SARS-CoV-2 virus propagation

Clinical isolates of SARS-CoV-2 variants were obtained from BEI resources: WA1/2020 (NR-52281), Alpha (NR-54011), Beta (NR-54009), Gamma (NR-54982), Epsilon (NR-55308) and (NR-55309), Delta (NR-55611). To propagate, a 70% confluent T25 flask of Vero E6 cells was infected at a MOI of 0.01 in diluted in 1 mL Opti-MEM for 1 hour at 37°C with occasional rocking. 4mL of D10 was then added and the flask was incubated for 72 hours at 37°C. Following incubation, flasks were checked for cytopathic effect (CPE), after which supernatant was collected and spun at 3000×g for 5 minutes to remove cellular debris, then aliquoted for storage at -80°C. Propagated stocks were titrated with 8 × 10-fold dilutions in a focus forming assay as described below.

### SARS-CoV-2 immunofluorescence

96-well TC plates were seeded to 50% confluency with VeroE6 or Caco2 cells. Plates were then inoculated at a MOI of 0.1 of SARS-CoV-2 in 50 μL of Opti-MEM for 1 hour at 37°C with occasional rocking. An additional 50 μL of fresh media was then added and incubated for 24 hours at 37°C. Plates were fixed by submerging in 4% para-formaldehyde (PFA) in PBS for 1 hour, then brought into BSL-1 for immunofluorescence staining. Permeabilization was performed with 2% bovine serum albumin (BSA), 0.1% Triton-X 100 in PBS for 1 hour at RT. Cells were stained with saRBD-1 at 2 μg/ml, mouse anti-dsRNA (Millipore sigma # MABE1134) 1:1000. Anti-mouse IgG AF555 (Abcam ab150114) or anti-llama IgM AF488-conjugated secondary antibodies were added at 1:500 dilution for 1 hour at RT (Invitrogen). Confocal imaging was performed with a Zeiss LSM 980 using a 63x Plan-Achromatic 1.4 NA oil immersion objective. Images were processed with Zeiss Zen Blue software. Maximum intensity *z*-projections were prepared in ImageJ. All antibody stain images were pseudocolored for visual consistency.

### Flow cytometry of S transfected cells

293T cells were seeded at 70% confluency on 24 well plates. Cells were then transfected with 1 μg of HDM_IDTSpike_fixK S plasmid as described above. 24 hours after transfection, cells were harvested by scraping and immediately stained with live-dead 405nm stain Zombie Violet (BioLegend). After live-dead staining, cells were washed 2x with PBS, then fixed with 4% PFA for 15 minutes at RT. Cells were then washed 2x with PBS, then stained with saRBD-1 VHH at 2 μg/ml, VHH52 at 2 μg/ml or phalloidin-Alexa Fluor 488 (AF488) (ThermoFisher Scientific A12379) for 1 hour at RT. Cells were washed 2x with PBS then treated with anti-VHH biotin (Jackson 128-065-232) secondary antibody (Invitrogen #A16061) was added at 1:500 dilution for 1 hour at RT. Cells were then washed 2x with PBS and treated with streptavidin-AF488 (Jackson 016-540-084) for 1 hour at RT. Cells were then resuspended in flow buffer (PBS, 3% FBS, 2mM EDTA), and filtered. Cells were then run on a BD FACSymphony flow cytometer with the assistance of the OHSU flow cytometry core.

### Thermal shift assay

Purified RBD and saRBD-1 protein were diluted to 10 µM in PBS. The combined RBD + saRBD-1 sample contained 10 µM of each protein. SYPRO Orange dye was obtained as a 5000× solution (ThermoFisher) and was diluted to a final concentration of 5× for all conditions. Experiments were performed on a StepOnePlus rtPCR system (Applied Biosystems) in a melting curve experiment with reporter set to ROX using a step and hold method starting at 25°C, ramping at 1°C per minute until 95°C in a total reaction volume of 50 µL per well. The melting point was calculated as the temperature with the minimum value of the first derivative of fluorescence emission as a function of temperature. The derivative values are calculated automatically by the StepOnePlus software and exported along with the normalized fluorescence intensity values.

### Spike-pseudotyped lentivirus production

293T cells were seeded at 2 million cells/dish in 6cm TC-treated dishes. The following day, cells were transfected as described above with lentivirus packaging plasmids, SARS-CoV-2 S plasmid, and LzGreen as described above (Crawford et al., 2020). After transfection, cells were incubated at 37C for 48 hours. Viral media was harvested, filtered with 0.45μm filter, then frozen before use. Virus transduction capability was titered on 293T-ACE2 cells in DMEM plus 5 µg/mL polybrene. LzGreen titers were determined by fluorescence using BZ-X700 all-in-one fluorescent microscope (Keyence): a 1:16 dilution was decided as optimal for following neutralization assays due to broad transduced foci distribution. Each well was captured by Keyence microscope and stitched using built-in software. GFP signal was quantified for the stitched images using Keyence software, transduction levels were normalized to virus only control wells.

### Enzyme-linked immunosorbent assay (ELISA)

MaxiSorp ELISA plates (Invitrogen), were coated with purified recombinant SARS-COV-2 RBD domain (BEI, NR-52306) at 2 μg/μl in PBS, or equivalent molar ratios of S1 domain (BEI, NR-53798), S2 domain (BEI, NR-53799), or trimer (BEI, NR-52396). Coating was carried out overnight at 4°C. Plates were blocked in wash buffer (2% BSA, 0.1% tween-20 in PBS) for 30 minutes at RT. Dilutions ranging from 14.2 nm to 3 pM of saRBD-1 or Fc-saRBD-1 were incubated for 1 hour at RT. Plates were washed with PBST (0.1% tween-20 in PBS) 4 times between each antibody addition. Anti-VHH -biotinylated antibody and streptavidin -HRP secondary antibodies were used at 1:10000 concentration in blocking buffer and were incubated 1 hour at RT. After the final wash, plates were incubated for 10 minutes with 50 μL of TMB HRP substrate (ThermoFisher) at RT, before adding 50 μL of stopping solution (2N H_2_SO_4_). Plate absorbances at 405nm were measured using a CLARIOstar® Plus plate fluorimeter (BMG Labtech).

### Biolayer interferometry (BLI)

Streptavidin biosensors (ForteBio) were soaked in PBS for at least 30 minutes prior to starting experiments. Biosensors were prepared with the following steps: equilibration in kinetics buffer (10 mM HEPES, 150 mM NaCl, 3mM EDTA, 0.005% Tween-20, 0.1% BSA, pH 7.5) for 300 seconds, loading of biotinylated RBD protein (10ug/mL) in kinetics buffer for 200 seconds, and blocking in 1 μM D-Biotin in kinetics buffer for 50 seconds. Binding was measured for seven 3-fold serial dilutions of each monoclonal antibody using the following cycle sequence: baseline for 300 seconds in kinetics buffer, association for 300 seconds with antibody diluted in kinetics buffer, dissociation for 750 seconds in plain kinetics buffer, and regeneration by 3 cycles of 20 seconds in 10 mM glycine pH 1.7, then 20 seconds in kinetics buffer. All antibodies were run against an isotype control antibody at the same concentration. For competition experiments, biosensors were loaded with RBD similarly to binding experiments, then bound with 50 nM saRBD-1 or 100 nM Fc-saRBD-1 for 300 seconds before transferring to monoclonal antibody diluted in kinetics buffer for 300 seconds (cov2rbdc2-mab10 was used at 20 nM, while srbdc4-mab10 and srbdc5-mab10 were used at 10 nM). Data analysis was performed using the ForteBio data analysis HT 10.0 software. Curves were reference subtracted using the isotype control and each cycle was aligned according to its baseline step. K_D_’s were calculated using a 1:1 binding model, and the kinetic parameters (K_D_, k_ON_, k_OFF_) were averaged from concentrations and replicates, excluding dilutions with an R_2_ less than 0.9 or an Rmax more than double the average of other concentrations.

### Pseudovirus neutralization assay

The neutralization protocol was based on previously reported neutralization methods utilizing SARS-CoV-2 S pseudotyped lentivirus (Crawford et al., 2020). 293T-ACE2 cells were seeded poly-lysine treated 96-well plates at a density of 10,000 cells per well. Cells were allowed to grow overnight at 37°C. LzGreen SARS-COV-2 S pseudotyped lentiviruses were mixed with saRBD-1, or VHH52 control antibody. Immunized alpaca serum was used as positive neutralization control, while virus alone was used as negative control. Dilutions of antibodies ranged from 177 nM to 170 pm for saRBD-1 and 26.3nM and 25 pM for Fc-saRBD-1, and 6.57 nM to 4 pM Bi-saRBD-1. Virus-antibody mixture was incubated at 37C for 1 hour after which polybrene was added up to 5 μg/ml and the mixture was added to 293T-ACE2 cells. Cells were incubated with neutralized virus for 44 hours before imaging. Cells were fixed with 4% PFA for 1 hour at RT. Fixed cells were washed with PBS 2x, then incubated with 10 μg/ml DAPI for 10 minutes at RT, imaged with BZ-X700 all-in-one fluorescent microscope (Keyence). Estimated area of DAPI and GFP fluorescent pixels were calculated with built in BZ-X software (Keyence).

### Focus forming assay (FFA)

The FFAs was performed as previously described (Case et al., 2020). In brief, Vero E6 cells were plated at 20,000 cells/well or Caco-2 cells were plated at 24,000 cells/well in 96-well plates and incubated overnight. Titrated SARS-CoV-2 stocks were diluted to 3,333 ffu/mL. To 20 μL of virus, 20 μL of antibody dilutions were added: saRBD-1, VHH52, or Fc-saRBD-1 were used at 8×4-fold serial dilutions ranging from 6.25 μg/mL to 381 pg/mL for saRBD-1, 1.25 μg/mL to 76.2 pg/mL for Fc-saRBD-1, and 420 μg/mL to 25.6 pg/mL All virus and antibody dilutions were prepared in Opti-MEM media. 30 μL of neutralized virus was then added to the confluent cells and incubated for 1 hour at 37°C. 150 μL of overlay media (Opti-MEM, 2% FBS, 2% Methylcellulose) was then added to each well and incubated for 24 hours at 37°C. Plates were fixed using 4% PFA for 1 hour at RT. Plates were then blocked for 30 minutes with permeabilization buffer (PBS, 0.1% BSA and 0.1% saponin). RBD immunized alpaca sera was used as a primary antibody at 1:5,000 dilution in permeabilization buffer, and anti-Llama-HRP secondary was used at 1:20,000 dilution in permeabilization buffer. Plates were developed in 30 µL TrueBlue (SeraCare) substrate and imaged with an Immunospot analyzer (CTL). Foci were counted with Viridot (Katzelnick et al., 2018) version 1.0 in R version 4.1.0.

### Quantification and statistical analysis

For ELISA and neutralization data, EC_50_ and IC_50_ values were calculated using python software pipeline based on input data. Curves were fit to each data set using the same pipeline. For ELISA data, EC_50_ were calculated from OD_450_ nm signal relative to maximal signal for a given pattern. Background was subtracted, then each was normalized to the maximum value for that antigen. The S2 domain data was analyzed differently, as it was comparable to background, background absorbance was first subtracted before normalization to maximum value.

For pseudovirus neutralization experiments, total surface area and intensity of blue and green signal were quantified using Keyence software. Three technical replicates were performed for each concentration in each experiment. The two values closest to the average of the triplicates were used to calculate the average of green signal (transduction), which was normalized to average blue signal (DAPI) for each concentration. The normalized transduction data were fit to a logistic function to determine EC_50_ and IC_50_ values in python version 3.8.10.

For live virus neutralization assays, focus counts generated with Viridot were manually checked for artifacts and recounted manually when incorrect. Focus counts were normalized in a plate-wise manner to the average of virus only well focus counts to obtain percent infection values, which were fit to a logistic function to determine FRNT_50_ values in python version 3.8.10.

## Acknowledgments

BLI data were generated on an Octet Red 384, which is made available and supported by OHSU Proteomics Shared Resource facility and equipment grant number S10OD023413. We also thank the OHSU Flow Cytometry Shared Resource, and OHSU Advanced Light Microscopy Core for the use of their software, equipment, and expertise.

## Data availability

The source data for this study are provided with the paper. SARS-CoV-2 RBD plasmid based on Wuhan isolate sequence from GenBank under accession number MN908947.3.

## Code availability

No unique code was generated as part of this study.

## Author Contributions

Conceptualization: F.G.T., J.B.W., and T.A.B.; methodology, formal analysis, and investigation: T.A.B., J.B.W., H.C.L., and S.K.M.; writing – original draft: J.B.W., T.A.B.; writing – review and editing: all authors; visualization: T.A.B., J.B.W., and F.G.T.; supervision: F.G.T.; project administration: F.G.T.; fund acquisition: F.G.T.

## Lead contact

Further information and requests for resources and reagents should be directed to the lead contact, Fikadu G. Tafesse

## References

1. Amanat, F., Nguyen, T., Chromikova, V., Strohmeier, S., Stadlbauer, D., Javier, A., Jiang, K., Asthagiri-Arunkumar, G., Polanco, J., Bermudez-Gonzalez, M., et al. (2020). A serological assay to detect SARS-CoV-2 seroconversion in humans (Allergy and Immunology).

2. Baden, L.R., El Sahly, H.M., Essink, B., Kotloff, K., Frey, S., Novak, R., Diemert, D., Spector, S.A., Rouphael, N., Creech, C.B., et al. (2021). Efficacy and Safety of the mRNA-1273 SARS-CoV-2 Vaccine. New England Journal of Medicine 384, 403–416.

3. Barnes, C.O., Jette, C.A., Abernathy, M.E., Dam, K.-M.A., Esswein, S.R., Gristick, H.B., Malyutin, A.G., Sharaf, N.G., Huey-Tubman, K.E., Lee, Y.E., et al. (2020). SARS-CoV-2 neutralizing antibody structures inform therapeutic strategies. Nature 588, 682–687.

4. Bates, T.A., Leier, H.C., Lyski, Z.L., McBride, S.K., Coulter, F.J., Weinstein, J.B., Goodman, J.R., Lu, Z., Siegel, S.A.R., Sullivan, P., et al. (2021a). Neutralization of SARS-CoV-2 variants by convalescent and BNT162b2 vaccinated serum. Nat Commun 12, 5135.

5. Bates, T.A., Leier, H.C., Lyski, Z.L., Goodman, J.R., Curlin, M.E., Messer, W.B., and Tafesse, F.G. (2021b). Age-Dependent Neutralization of SARS-CoV-2 and P.1 Variant by Vaccine Immune Serum Samples. JAMA 326, 868–869.

6. Bates, T.A., Weinstein, J.B., Farley, S., Leier, H.C., Messer, W.B., and Tafesse, F.G. (2021c). Cross-reactivity of SARS-CoV structural protein antibodies against SARS-CoV-2. Cell Reports 34, 108737.

7. Bates, T.A., McBride, S.K., Winders, B., Schoen, D., Trautmann, L., Curlin, M.E., and Tafesse, F.G. (2022). Antibody Response and Variant Cross-Neutralization After SARS-CoV-2 Breakthrough Infection. JAMA 327, 179–181.

8. Bierig, T., Collu, G., Blanc, A., Poghosyan, E., and Benoit, R.M. (2020). Design, Expression, Purification, and Characterization of a YFP-Tagged 2019-nCoV Spike Receptor-Binding Domain Construct. Front. Bioeng. Biotechnol. 0.

9. Carrillo, J., Izquierdo-Useros, N., Ávila-Nieto, C., Pradenas, E., Clotet, B., and Blanco, J. (2021). Humoral immune responses and neutralizing antibodies against SARS-CoV-2; implications in pathogenesis and protective immunity. Biochem Biophys Res Commun 538, 187–191.

10. Case, J.B., Rothlauf, P.W., Chen, R.E., Liu, Z., Zhao, H., Kim, A.S., Bloyet, L.-M., Zeng, Q., Tahan, S., Droit, L., et al. (2020). Neutralizing Antibody and Soluble ACE2 Inhibition of a Replication-Competent VSV-SARS-CoV-2 and a Clinical Isolate of SARS-CoV-2. Cell Host Microbe 28, 475–485.e5.

11. Cavallari, M. (2017). Rapid and Direct VHH and Target Identification by Staphylococcal Surface Display Libraries. Int J Mol Sci 18.

12. Chen, R.E., Zhang, X., Case, J.B., Winkler, E.S., Liu, Y., VanBlargan, L.A., Liu, J., Errico, J.M., Xie, X., Suryadevara, N., et al. (2021). Resistance of SARS-CoV-2 variants to neutralization by monoclonal and serum-derived polyclonal antibodies. Nature Medicine 1–10.

13. Crawford, K.H.D., Eguia, R., Dingens, A.S., Loes, A.N., Malone, K.D., Wolf, C.R., Chu, H.Y., Tortorici, M.A., Veesler, D., Murphy, M., et al. (2020). Protocol and reagents for pseudotyping lentiviral particles with SARS-CoV-2 Spike protein for neutralization assays (Microbiology).

14. Crommelin, D.J.A., Anchordoquy, T.J., Volkin, D.B., Jiskoot, W., and Mastrobattista, E. (2021). Addressing the Cold Reality of mRNA Vaccine Stability. Journal of Pharmaceutical Sciences 110, 997–1001.

15. Daniel, W., Nivet, M., Warner, J., and Podolsky, D.K. (2021). Early Evidence of the Effect of SARS-CoV-2 Vaccine at One Medical Center. New England Journal of Medicine 384, 1962– 1963.

16. Deng, X., Garcia-Knight, M.A., Khalid, M.M., Servellita, V., Wang, C., Morris, M.K., Sotomayor-González, A., Glasner, D.R., Reyes, K.R., Gliwa, A.S., et al. (2021). Transmission, infectivity, and neutralization of a spike L452R SARS-CoV-2 variant. Cell 184, 3426–3437.e8.

17. Dong, E., Du, H., and Gardner, L. (2020). An interactive web-based dashboard to track COVID-19 in real time. The Lancet Infectious Diseases 20, 533–534.

18. Eppendorf, N.S. Faster Isolation of PBMC Using Ficoll-Paque® Plus in the Eppendorf Multipurpose Benchtop Centrifuges 5920 R and 5910 Ri. 6.

19. Eroshenko, N., Gill, T., Keaveney, M.K., Church, G.M., Trevejo, J.M., and Rajaniemi, H. (2020). Implications of antibody-dependent enhancement of infection for SARS-CoV-2 countermeasures. Nat Biotechnol 38, 789–791.

20. Frei, J.C., and Lai, J.R. (2016). Protein and Antibody Engineering by Phage Display. Methods Enzymol 580, 45–87.

21. Glasgow, A., Glasgow, J., Limonta, D., Solomon, P., Lui, I., Zhang, Y., Nix, M.A., Rettko, N.J., Zha, S., Yamin, R., et al. (2020). Engineered ACE2 receptor traps potently neutralize SARS-CoV-2. PNAS 117, 28046–28055.

22. Gordon, D.E., Jang, G.M., Bouhaddou, M., Xu, J., Obernier, K., O’Meara, M.J., Guo, J.Z., Swaney, D.L., Tummino, T.A., Huttenhain, R., et al. (2020). A SARS-CoV-2-Human Protein-Protein Interaction Map Reveals Drug Targets and Potential Drug-Repurposing (Systems Biology).

23. Greaney, A.J., Starr, T.N., Gilchuk, P., Zost, S.J., Binshtein, E., Loes, A.N., Hilton, S.K., Huddleston, J., Eguia, R., Crawford, K.H.D., et al. (2021). Complete Mapping of Mutations to the SARS-CoV-2 Spike Receptor-Binding Domain that Escape Antibody Recognition. Cell Host & Microbe 29, 44–57.e9.

24. Günaydın, G., Yu, S., Gräslund, T., Hammarström, L., and Marcotte, H. (2016). Fusion of the mouse IgG1 Fc domain to the VHH fragment (ARP1) enhances protection in a mouse model of rotavirus. Scientific Reports 6, 30171.

25. Güttler, T., Aksu, M., Dickmanns, A., Stegmann, K.M., Gregor, K., Rees, R., Taxer, W., Rymarenko, O., Schünemann, J., Dienemann, C., et al. (2021). Neutralization of SARS-CoV-2 by highly potent, hyperthermostable, and mutation-tolerant nanobodies. EMBO J 40, e107985.

26. Hanke, L., Vidakovics Perez, L., Sheward, D.J., Das, H., Schulte, T., Moliner-Morro, A., Corcoran, M., Achour, A., Karlsson Hedestam, G.B., Hällberg, B.M., et al. (2020). An alpaca nanobody neutralizes SARS-CoV-2 by blocking receptor interaction. Nature Communications 11, 4420.

27. Hanke, L., Das, H., Sheward, D.J., Perez Vidakovics, L., Urgard, E., Moliner-Morro, A., Kim, C., Karl, V., Pankow, A., Smith, N.L., et al. (2022). A bispecific monomeric nanobody induces spike trimer dimers and neutralizes SARS-CoV-2 in vivo. Nat Commun 13, 155.

28. Hoffmann, M., Kleine-Weber, H., Schroeder, S., Krüger, N., Herrler, T., Erichsen, S., Schiergens, T.S., Herrler, G., Wu, N.-H., Nitsche, A., et al. (2020). SARS-CoV-2 Cell Entry Depends on ACE2 and TMPRSS2 and Is Blocked by a Clinically Proven Protease Inhibitor. Cell 181, 271–280.e8.

29. Hoffmann, M., Arora, P., Groß, R., Seidel, A., Hörnich, B.F., Hahn, A.S., Krüger, N., Graichen, L., Hofmann-Winkler, H., Kempf, A., et al. (2021). SARS-CoV-2 variants B.1.351 and P.1 escape from neutralizing antibodies. Cell 184, 2384–2393.e12.

30. Holder, J. Tracking Coronavirus Vaccinations Around the World. The New York Times.

31. Hu, X., Hu, C., Jiang, D., Zuo, Q., Li, Y., Wang, Y., and Chen, X. (2020). Effectiveness of Convalescent Plasma Therapy for COVID-19 Patients in Hunan, China. Dose-Response 18, 1559325820979921.

32. Huang, C., Wang, Y., Li, X., Ren, L., Zhao, J., Hu, Y., Zhang, L., Fan, G., Xu, J., Gu, X., et al. (2020). Clinical features of patients infected with 2019 novel coronavirus in Wuhan, China. The Lancet 395, 497–506.

33. Ingram, J.R., Schmidt, F.I., and Ploegh, H.L. (2018). Exploiting Nanobodies’ Singular Traits. Annual Review of Immunology 36, 695–715.

34. Jiang, S., Hillyer, C., and Du, L. (2020). Neutralizing Antibodies against SARS-CoV-2 and Other Human Coronaviruses. Trends Immunol 41, 355–359.

35. Jones, B.E., Brown-Augsburger, P.L., Corbett, K.S., Westendorf, K., Davies, J., Cujec, T.P., Wiethoff, C.M., Blackbourne, J.L., Heinz, B.A., Foster, D., et al. (2021). The neutralizing antibody, LY-CoV555, protects against SARS-CoV-2 infection in nonhuman primates. Science Translational Medicine.

36. Kartoglu, U.H., Moore, K.L., and Lloyd, J.S. (2020). Logistical challenges for potential SARS-CoV-2 vaccine and a call to research institutions, developers and manufacturers. Vaccine 38, 5393–5395.

37. Katzelnick, L.C., Escoto, A.C., McElvany, B.D., Chávez, C., Salje, H., Luo, W., Rodriguez-Barraquer, I., Jarman, R., Durbin, A.P., Diehl, S.A., et al. (2018). Viridot: An automated virus plaque (immunofocus) counter for the measurement of serological neutralizing responses with application to dengue virus. PLOS Neglected Tropical Diseases 12, e0006862.

38. Khoury, D.S., Cromer, D., Reynaldi, A., Schlub, T.E., Wheatley, A.K., Juno, J.A., Subbarao, K., Kent, S.J., Triccas, J.A., and Davenport, M.P. (2021). Neutralizing antibody levels are highly predictive of immune protection from symptomatic SARS-CoV-2 infection. Nat Med 1–7.

39. Kim, C., Ryu, D.-K., Lee, J., Kim, Y.-I., Seo, J.-M., Kim, Y.-G., Jeong, J.-H., Kim, M., Kim, J.-I., Kim, P., et al. (2021). A therapeutic neutralizing antibody targeting receptor binding domain of SARS-CoV-2 spike protein. Nature Communications 12, 288.

40. Koenig, P.-A., Das, H., Liu, H., Kümmerer, B.M., Gohr, F.N., Jenster, L.-M., Schiffelers, L.D.J., Tesfamariam, Y.M., Uchima, M., Wuerth, J.D., et al. (2021). Structure-guided multivalent nanobodies block SARS-CoV-2 infection and suppress mutational escape. Science.

41. Korber, B., Fischer, W.M., Gnanakaran, S., Yoon, H., Theiler, J., Abfalterer, W., Hengartner, N., Giorgi, E.E., Bhattacharya, T., Foley, B., et al. (2020). Tracking Changes in SARS-CoV-2 Spike: Evidence that D614G Increases Infectivity of the COVID-19 Virus. Cell 182, 812–827.e19.

42. Kumar, S., Chandele, A., and Sharma, A. (2021). Current status of therapeutic monoclonal antibodies against SARS-CoV-2. PLOS Pathogens 17, e1009885.

43. Laursen, N.S., Friesen, R.H.E., Zhu, X., Jongeneelen, M., Blokland, S., Vermond, J., Eijgen, A. van, Tang, C., Diepen, H. van, Obmolova, G., et al. (2018). Universal protection against influenza infection by a multidomain antibody to influenza hemagglutinin. Science 362, 598–602.

44. Levin, E.G., Lustig, Y., Cohen, C., Fluss, R., Indenbaum, V., Amit, S., Doolman, R., Asraf, K., Mendelson, E., Ziv, A., et al. (2021). Waning Immune Humoral Response to BNT162b2 Covid-19 Vaccine over 6 Months. New England Journal of Medicine 0, null.

45. Liu, H., Zhang, Q., Wei, P., Chen, Z., Aviszus, K., Yang, J., Downing, W., Jiang, C., Liang, B., Reynoso, L., et al. (2021a). The basis of a more contagious 501Y.V1 variant of SARS-CoV-2. Cell Res 31, 720–722.

46. Liu, J., Liu, Y., Xia, H., Zou, J., Weaver, S.C., Swanson, K.A., Cai, H., Cutler, M., Cooper, D., Muik, A., et al. (2021b). BNT162b2-elicited neutralization of B.1.617 and other SARS-CoV-2 variants. Nature 1–5.

47. Liu, L., Iketani, S., Guo, Y., Chan, J.F.-W., Wang, M., Liu, L., Luo, Y., Chu, H., Huang, Y., Nair, M.S., et al. (2021c). Striking Antibody Evasion Manifested by the Omicron Variant of SARS-CoV-2. Nature 1–8.

48. Luo, R., Delaunay-Moisan, A., Timmis, K., and Danchin, A. (2021). SARS-CoV-2 biology and variants: anticipation of viral evolution and what needs to be done. Environmental Microbiology 23, 2339–2363.

49. Maass, D.R., Sepulveda, J., Pernthaner, A., and Shoemaker, C.B. (2007). Alpaca (Lama pacos) as a convenient source of recombinant camelid heavy chain antibodies (VHHs). Journal of Immunological Methods 324, 13–25.

50. Mast, F.D., Fridy, P.C., Ketaren, N.E., Wang, J., Jacobs, E.Y., Olivier, J.P., Sanyal, T., Molloy, K.R., Schmidt, F., Rutkowska, M., et al. (2021). Highly synergistic combinations of nanobodies that target SARS-CoV-2 and are resistant to escape. ELife 10, e73027.

51. Mathieu, E., Ritchie, H., Ortiz-Ospina, E., Roser, M., Hasell, J., Appel, C., Giattino, C., and Rodés-Guirao, L. (2021). A global database of COVID-19 vaccinations. Nat Hum Behav 1–7.

52. Pinto, D., Park, Y.-J., Beltramello, M., Walls, A.C., Tortorici, M.A., Bianchi, S., Jaconi, S., Culap, K., Zatta, F., De Marco, A., et al. (2020). Cross-neutralization of SARS-CoV-2 by a human monoclonal SARS-CoV antibody. Nature 583, 290–295.

53. Planas, D., Bruel, T., Grzelak, L., Guivel-Benhassine, F., Staropoli, I., Porrot, F., Planchais, C., Buchrieser, J., Rajah, M.M., Bishop, E., et al. (2021). Sensitivity of infectious SARS-CoV-2 B.1.1.7 and B.1.351 variants to neutralizing antibodies. Nat Med 27, 917–924.

54. Pymm, P., Adair, A., Chan, L.-J., Cooney, J.P., Mordant, F.L., Allison, C.C., Lopez, E., Haycroft, E.R., O’Neill, M.T., Tan, L.L., et al. (2021). Nanobody cocktails potently neutralize SARS-CoV-2 D614G N501Y variant and protect mice. Proc Natl Acad Sci U S A 118.

55. Rotman, M., Welling, M.M., van den Boogaard, M.L., Moursel, L.G., van der Graaf, L.M., van Buchem, M.A., van der Maarel, S.M., and van der Weerd, L. (2015). Fusion of hIgG1-Fc to 111In-anti-amyloid single domain antibody fragment VHH-pa2H prolongs blood residential time in APP/PS1 mice but does not increase brain uptake. Nuclear Medicine and Biology 42, 695– 702.

56. Saerens, D., Kinne, J., Bosmans, E., Wernery, U., Muyldermans, S., and Conrath, K. (2004). Single Domain Antibodies Derived from Dromedary Lymph Node and Peripheral Blood Lymphocytes Sensing Conformational Variants of Prostate-specific Antigen. J. Biol. Chem. 279, 51965–51972.

57. Salvador, J.-P., Vilaplana, L., and Marco, M.-P. (2019). Nanobody: outstanding features for diagnostic and therapeutic applications. Anal Bioanal Chem 411, 1703–1713.

58. Schmidt, F.I. (2018). Phenotypic Lentivirus Screens to Identify Antiviral Single Domain Antibodies. In Influenza Virus: Methods and Protocols, Y. Yamauchi, ed. (New York, NY: Springer New York), pp. 139–158.

59. Schoof, M., Faust, B., Saunders, R.A., Sangwan, S., Rezelj, V., Hoppe, N., Boone, M., Billesbølle, C.B., Puchades, C., Azumaya, C.M., et al. (2020). An ultrapotent synthetic nanobody neutralizes SARS-CoV-2 by stabilizing inactive Spike. Science 370, 1473–1479.

60. Shan, D., Press, O.W., Tsu, T.T., Hayden, M.S., and Ledbetter, J.A. (1999). Characterization of scFv-Ig Constructs Generated from the Anti-CD20 mAb 1F5 Using Linker Peptides of Varying Lengths. The Journal of Immunology 162, 6589–6595.

61. Shen, X., Tang, H., Pajon, R., Smith, G., Glenn, G.M., Shi, W., Korber, B., and Montefiori, D.C. (2021). Neutralization of SARS-CoV-2 Variants B.1.429 and B.1.351. New England Journal of Medicine 0, null.

62. Shrotri, M., Navaratnam, A.M.D., Nguyen, V., Byrne, T., Geismar, C., Fragaszy, E., Beale, S., Fong, W.L.E., Patel, P., Kovar, J., et al. (2021). Spike-antibody waning after second dose of BNT162b2 or ChAdOx1. Lancet.

63. Tiller, T., Meffre, E., Yurasov, S., Tsuiji, M., Nussenzweig, M.C., and Wardemann, H. (2008). Efficient generation of monoclonal antibodies from single human B cells by single cell RT-PCR and expression vector cloning. Journal of Immunological Methods 329, 112–124.

64. Wagner, T.R., Schnepf, D., Beer, J., Ruetalo, N., Klingel, K., Kaiser, P.D., Junker, D., Sauter, M., Traenkle, B., Frecot, D.I., et al. (2021). Biparatopic nanobodies protect mice from lethal challenge with SARS-CoV-2 variants of concern. EMBO Rep e53865.

65. Weinreich, D.M., Sivapalasingam, S., Norton, T., Ali, S., Gao, H., Bhore, R., Musser, B.J., Soo, Y., Rofail, D., Im, J., et al. (2021). REGN-COV2, a Neutralizing Antibody Cocktail, in Outpatients with Covid-19. New England Journal of Medicine 384, 238–251.

66. Wrapp, D., De Vlieger, D., Corbett, K.S., Torres, G.M., Wang, N., Van Breedam, W., Roose, K., van Schie, L., Hoffmann, M., Pöhlmann, S., et al. (2020a). Structural Basis for Potent Neutralization of Betacoronaviruses by Single-Domain Camelid Antibodies. Cell 181, 1004–1015.e15.

67. Wrapp, D., Vlieger, D.D., Corbett, K.S., Torres, G.M., Breedam, W.V., Roose, K., Schie, L. van, Team, V.-C.C.-19 R., Hoffmann, M., Pöhlmann, S., et al. (2020b). Structural Basis for Potent Neutralization of Betacoronaviruses by Single-domain Camelid Antibodies. BioRxiv 2020.03.26.010165.

68. Wu, Y., Wang, F., Shen, C., Peng, W., Li, D., Zhao, C., Li, Z., Li, S., Bi, Y., Yang, Y., et al. (2020). A noncompeting pair of human neutralizing antibodies block COVID-19 virus binding to its receptor ACE2. Science 368, 1274–1278.

69. Xiang, Y., Nambulli, S., Xiao, Z., Liu, H., Sang, Z., Duprex, W.P., Schneidman-Duhovny, D., Zhang, C., and Shi, Y. (2020). Versatile and multivalent nanobodies efficiently neutralize SARS-CoV-2. Science 370, 1479–1484.

70. Xu, J., Xu, K., Jung, S., Conte, A., Lieberman, J., Muecksch, F., Lorenzi, J.C.C., Park, S., Schmidt, F., Wang, Z., et al. (2021). Nanobodies from camelid mice and llamas neutralize SARS-CoV-2 variants. Nature.

71. Yuan, M., Wu, N.C., Zhu, X., Lee, C.-C.D., So, R.T.Y., Lv, H., Mok, C.K.P., and Wilson, I.A. (2020). A highly conserved cryptic epitope in the receptor binding domains of SARS-CoV-2 and SARS-CoV. Science 368, 630–633.

72. Zeng, H., Wang, D., Nie, J., Liang, H., Gu, J., Zhao, A., Xu, L., Lang, C., Cui, X., Guo, X., et al. (2020). The efficacy assessment of convalescent plasma therapy for COVID-19 patients: a multi-center case series. Sig Transduct Target Ther 5, 1–12.

73. Zhang, X., Wu, S., Wu, B., Yang, Q., Chen, A., Li, Y., Zhang, Y., Pan, T., Zhang, H., and He, X. (2021). SARS-CoV-2 Omicron strain exhibits potent capabilities for immune evasion and viral entrance. Sig Transduct Target Ther 6, 1–3.

74. Zupancic, J.M., Schardt, J.S., Desai, A.A., Makowski, E.K., Smith, M.D., Pornnoppadol, G., Garcia de Mattos Barbosa, M., Cascalho, M., Lanigan, T.M., and Tessier, P.M. (2021). Engineered Multivalent Nanobodies Potently and Broadly Neutralize SARS-CoV-2 Variants. Advanced Therapeutics 4, 2100099.

