## Supplemental Figures for "A potent alpaca-derived nanobody that neutralizes SARS-CoV-2 variants"

**A**

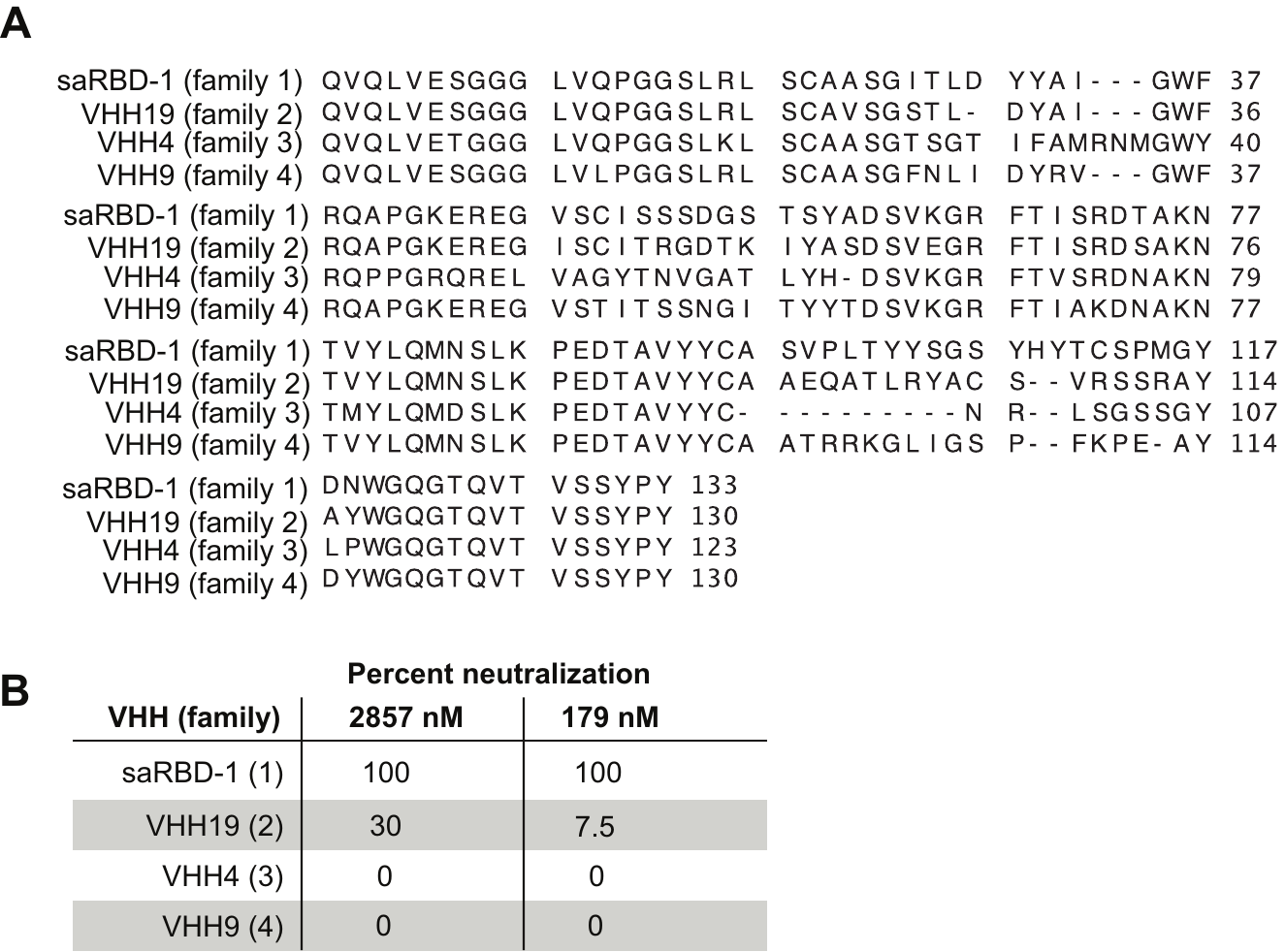

Figure S1: Neutralization capabilities of the dominant families of the anti-RBD VHH hits. A) Percent neutralization of S pseudotyped lentivirus by representative VHHs from 4 dominant families shown, including saRBD-1, at concentrations of 2857 nM and 179 nM. Signal normalized against virus-only control. Data presented from two independent experiments performed in triplicates.

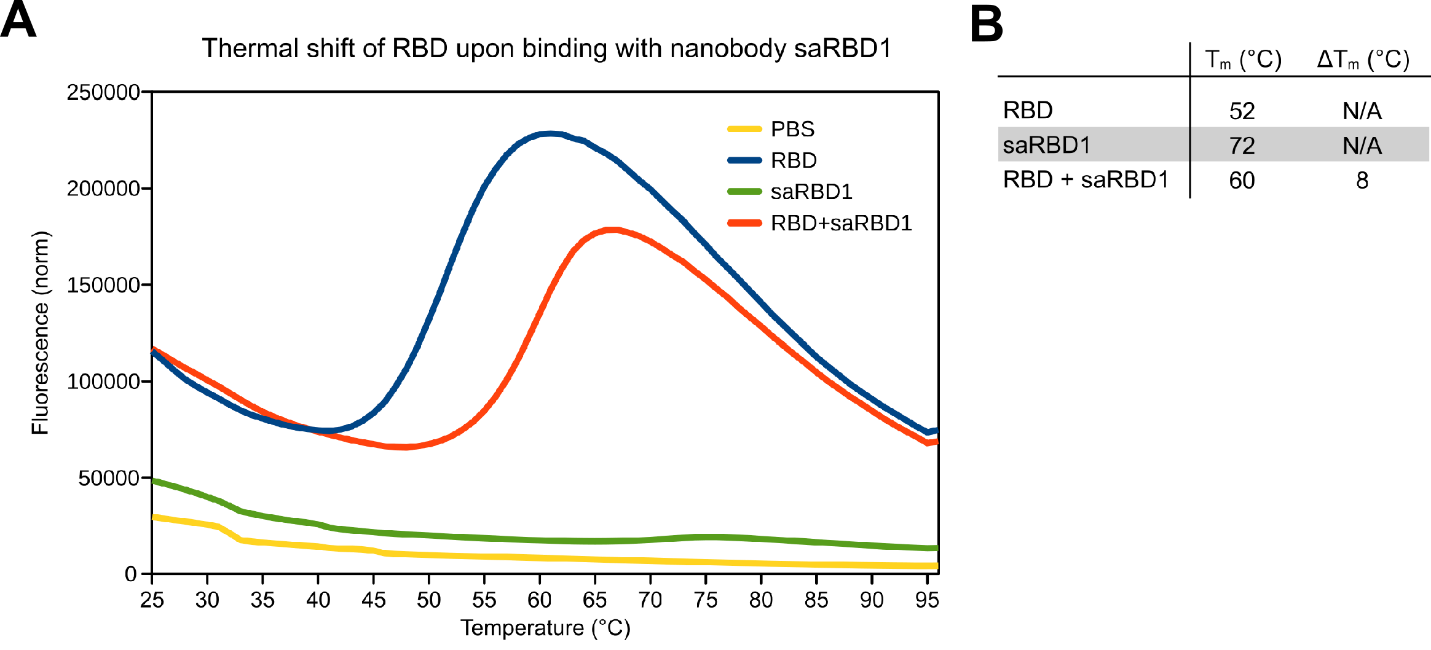

Figure S2: saRBD-1 stabilizes RBD in a thermal shift assay. A) Representative melting curves showing fluorescence intensity versus temperature in thermal shift experiments with Purified RBD protein diluted to 10µM in PBS is stabilized by addition of 10µM saRBD-1. B) Melting temperatures of RBD and saRBD-1 separately and combined. Melting temperature values are the minimum value of the first derivative of normalized fluorescence intensity with respect to temperature and the values shown are the average of three replicates.

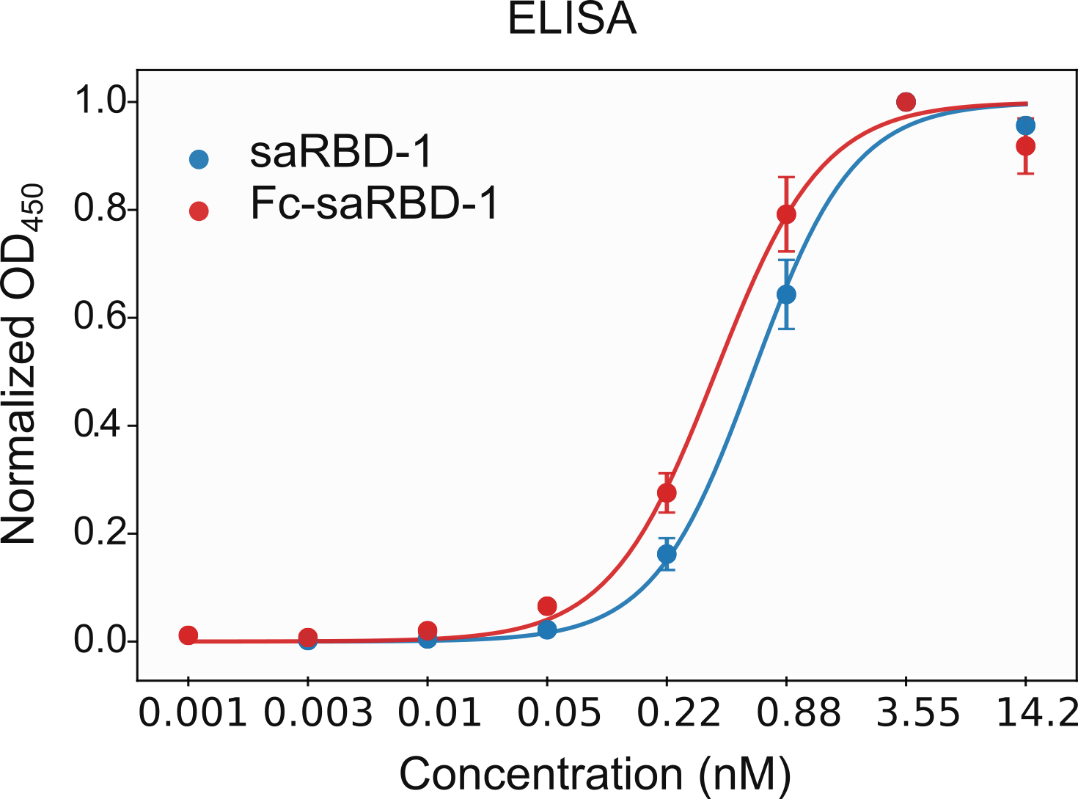

Figure S3: ELISA EC_50_ measurements of saRBD-1 and Fc-saRBD-1. Plates were coated with SARS-CoV-2 RBD protein. The EC50 of saRBD-1 is 607 pM and of Fc-saRBD-1 is 393 pM. Curves show the average of 3 replicate experiments and error bars represent standard error.
